# Transcriptional and epigenetic characterization of early striosomes identifies Foxf2 and Olig2 as factors required for development of striatal compartmentation and neuronal phenotypic differentiation

**DOI:** 10.1101/2020.05.19.105171

**Authors:** Maria-Daniela Cirnaru, Sicheng Song, Kizito-Tshitoko Tshilenge, Chuhyon Corwin, Justyna Mleczko, Carlos Galicia Aguirre, Houda Benlhabib, Jaroslav Bendl, Pasha Apontes, John F. Fullard, Jordi Creus Muncunill, Azadeh Reyah, Ali M. Nik, Peter Carlsson, Panos Roussos, Sean D. Mooney, Lisa M. Ellerby, Michelle E. Ehrlich

**Affiliations:** Department of Neurology, Icahn School of Medicine at Mount Sinai, New York, New York 10029, USA; Department of Epidemiology, University of Washington, Seattle, WA, USA; Buck Institute for Research on Aging, Novato, CA, 94945, USA; Pamela Sklar Division of Psychiatric Genomics, Icahn School of Medicine at Mount Sinai, One Gustave L. Levy Place, New York, NY, 10029, USA; Institute for Genomics and Multiscale Biology, Department of Genetics and Genomic Sciences, Icahn School of Medicine at Mount Sinai, One Gustave L. Levy Place, New York, NY, 10029, USA; Department of Psychiatry, Icahn School of Medicine at Mount Sinai, One Gustave L. Levy Place, New York, NY, 10029, USA; Department of Chemistry and Molecular Biology, University of Gothenburg, Gothenburg, Sweden; Mental Illness Research, Education, and Clinical Center (VISN 2 South), James J. Peters VA Medical Center, Bronx, NY 10468, USA

## Abstract

The basal ganglia, best known for processing information required for multiple aspects of movement, is also part of a network which regulates reward and cognition. The major output nucleus of the basal ganglia is the striatum, and its functions are dependent on neuronal compartmentation, including striosomes and matrix, which are selectively affected in disease. Striatal projection neurons are GABAergic medium spiny neurons (MSNs), all of which share basic molecular signatures but are subtyped by selective expression of receptors, neuropeptides, and other gene families. Neurogenesis of the striosome and matrix occurs in separate waves, but the factors regulating terminal neuronal differentiation following migration are largely unidentified. We performed RNA- and ATAC-seq on sorted murine striosome and matrix cells at postnatal day 3. Focusing on the striosomal compartment, we validated the localization and role of transcription factors and their regulator(s), previously not known to be associated with striatal development, including *Irx1, Foxf2, Olig2* and *Stat1/2*. In addition, we validated the enhancer function of a striosome-specific open chromatin region located 15Kb downstream of the *Olig2* gene. These data and data bases provide novel tools to dissect and manipulate the networks regulating MSN compartmentation and differentiation and thus provide new approaches to establishing MSN subtypes from human iPSCs for disease modeling and drug discovery.

## Introduction

Multiple diseases including Huntington’s (HD), Parkinson’s, X-linked dystonia-parkinsonism (XDP), addiction, autism, and schizophrenia are linked to dysregulation of the striatum. The dorsal striatum is comprised of the caudate and putamen in humans but consists of a single nucleus in the mouse. This nucleus is a key component of cortical and subcortical circuits regulating movement, reward, and aspects of cognition, including speech and language. The striatum is biochemically separated into two main compartments known as striosomes (patch) and matrix. Striosomes represent approximately 10-15 percent of the volume and are dispersed throughout the 80-85 percent occupied by the matrix. Importantly, imbalance between striatal compartments likely contributes to movement disorders^1-3^ and compartmentation also appears to be required for non-motor functions, e.g. speech and language ^4-9^. Notably, XDP appears to be due to preferential degeneration of the striosomes^10,11^ and there is possibly early, preferential loss of striosomal neurons in HD^12,13^. Currently, a suitable differentiation protocol for the generation of striosomal cells for *in vitro* studies and replacement therapies does not exist^14-19^.

Medium spiny projection neurons (MSNs) account for 85-95% of striatal neurons and are its only output neuron. MSNs receive massive glutamatergic and dopaminergic input, and project to the substantia nigra, globus pallidus, and thalamus, with minor projections to other regions. MSNs are morphologically homogeneous, but they are phenotypically heterogeneous and adult subtypes may be distinguished by unique transcriptomes^20,21^. The two major classifications of MSNs are direct vs. indirect, among which they are equally distributed^22^. Direct neurons (dMSNs) express the dopamine D1 receptor (D1R) and project directly to the substantia nigra (SN) or to the internal segment of the globus pallidus. The indirect neurons (iMSNs) express the dopamine D2 and adenosine 2A receptors (D2R, A2aR) and project to the external segment of the globus pallidus or to the subthalamic nucleus, which projects to the SN.

The distinction between striosome and matrix is based on differences in gene expression, by the origins of afferents from cortical regions, e.g. sensorimotor, limbic, and associative, and to some extent by the destination of their efferents^2,23,24^. They both contain direct and indirect neurons, and the striosome content of dMSNs appears to vary with region^22,25^. Amongst the most commonly used phenotypic markers in the adult are the striosomal mu opiate receptor (MOR) encoded by *Oprm1*, and the matrix calbindin, encoded by *Calb1*^2^. Striosome and matrix neurons also differ in their electrophysiologic properties^26^ and further mediate functions based on their connectivity ^27^. The striosome compartment has been specifically associated with stimulus-response learning^28^ and perhaps stereotypy^29,30^. Most recently, striosome neurons were identified as the major contributors to the nigrostriatal tract arising from dMSNs^27^.

There are multiple identified transcription factors (TFs) that are required for striatal induction and differentiation^14,18,31-40^, particularly into dMSNs and iMSNs. Some of these TFs were used for direct conversion protocols of human fibroblasts into MSNs^18,19^ and importantly, a widely used protocol yielded only a matrix phenotype ^18^, and several yielded largely Calb1+ neurons, with no mention of a striosome phenotype ^14-17,41^.

Replication of striatal development and function *in vitro* requires communication between compartments ^26^, and only a handful of TFs and signaling systems have been shown to define striosome/matrix compartmentation, e.g. *Ikaros-1*, retinoids, and *Nr4a1*^22,36,37,42-45^. We sought to use an unbiased transcriptomic and epigenetic approach, i.e. RNA seq and ATAC-seq, to identify TFs and open chromatin regions (OCRs) associated with striosome maturation specifically after they are first formed. This time point may also include terminal differentiation effectors as defined by Hobert^46^ which initiate and maintain the adult identity of neurons. To this end, we used the GENSAT *Nr4a1*-EGFP mouse, in which reporter expression is directed to striosomes throughout the life of the mouse^22,47^, and focused on striosome/matrix distinctions at postnatal day three (PND3). We opted to perform bulk RNA-seq rather than single cell to ensure the capture of low expression genes. As the complementary results to our analyses of EGFP-positive striosomal neurons, we also developed highly useful databases of the EGFP-negative, developing matrix neurons. The identified TFs and OCRs greatly add to the understanding of MSN development and allow for the generation of MSN subtypes for disease modeling.

## Results

### Transcriptional analysis of *Nr4a1*-GFP mice identify striatal compartment specific transcription factors *in vivo*

The transcription factor *Nr4a1* is a member of the Nur family of steroid/thyroid-like receptors^48^ and is expressed in the mouse striosome MSNs as early as embryonic day 14.5^47^. In this study, GENSAT transgenic mice expressing the reporter EGFP from a transgenic *Nr4a1* bacterial artificial chromosome (*Nr4a1*-EGFP) were used at PND3 (Fig. 1a). Fluorescence-activated cell sorting (FACS) was used to isolate the *Nr4a1*-EGFP positive and negative neurons representing striosome and matrix MSN populations, respectively (Fig. 1b). Visual inspection confirmed the virtual absence of EGFP^+^ neurons in the EGFP^−^ population, and vice versa (Fig. 1c). It is important to note that the EGFP^+^ cells also contained isolated, *Nr4a1*^*+*^ neurons that exist outside of discrete striosomes, and other non-neuronal *Nr4a1*^*+*^ cell types^49-52^. It is unknown whether the extra-striosomal neurons represent the exo-patch.

**Fig. 1:**
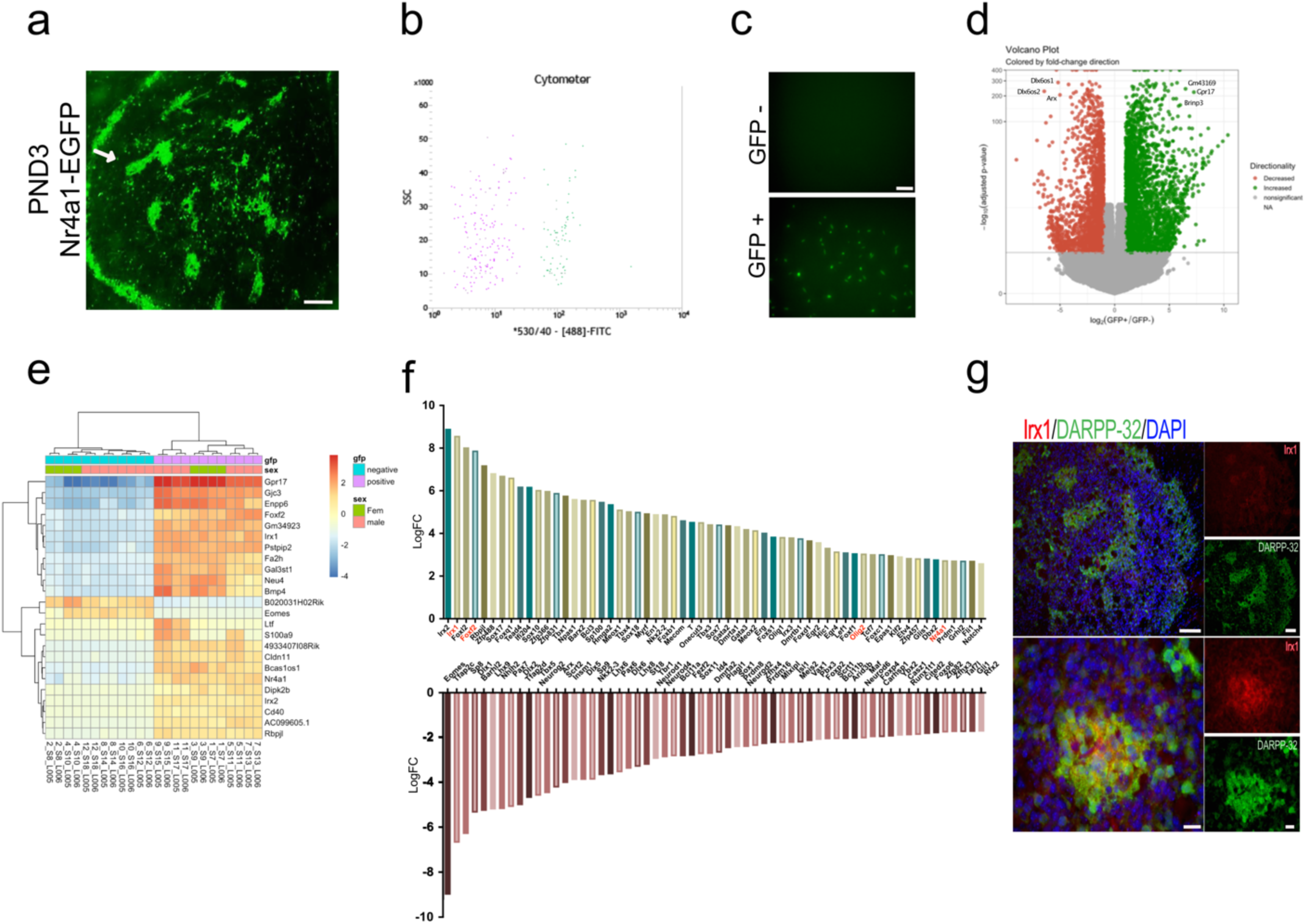
Identification of striatal striosome and matrix compartment-specific transcription factors using RNA-seq analysis in FACs EGFP^+^ and EGFP^−^ cells populations from the striata of PND3 Nr4a1-GFP mice. **a)** Coronal section of PND3 *Nr4a1*-EGFP mouse showing the striosome localization of EGFP indicated by the arrow. Scale bar (100 µm). **b)** Representative FACs cytometer panel indicating cell separation following the intensity of 488-FITC channel with clear distribution of the EGFP^+^ population on the right side of the panel. **c)** Detection of spontaneous GFP fluorescence of EGFP^+^ and EGFP^−^ cells seeded at 1000 cells/well 2 h after FACs sorting indicating the presence of EGFP exclusively in EGFP^+^ samples. Scale bar corresponds to 50 µm. **d)** Volcano plot visualizing the difference between the EGFP^+^ and EGFP^−^ gene expression. Genes with an adjusted p-value below 0.01 with absolute log2 fold ratio greater than 1 are highlighted. In red, the genes are relatively decreased in expression in EGFP^+^ population, i.e. enriched in the EGFP^−^ population, and in green, the genes that are relatively increased in expression in the EGFP^+^ population, and in grey, those genes that are equally distributed among the two populations. **e)** Heatmap of relative normalized counts across samples reporting of genes with their absolute log2 fold change greater than seven, and mean of their normalized counts of all samples greater than 40. **f)** Representation of the top 60 transcription factors enriched according to their log2 fold change in the EGFP^+^ population (upper panel) and in the EGFP^−^ population (lower panel). **g)** Representative low and high-power micrographs of Irx1 (red), DARPP-32 (green) and DAPI immunolabelling on 16 ⍰ m thick coronal sections from wild type PND3 showing the localization of Irx1 within the DARPP-32 immunopositive striosomes. Scale bars correspond to 100 µm and 20 µm respectively.

Next, we carried out transcriptomic analysis (RNA-seq) on both the EGFP^+^ and EGFP^−^ cells (Table 1). A total of 9124 genes were differentially expressed in the two compartments, with 4714 enriched in the EGFP^+^ cells (positive log2FC) and 4410 enriched in the EGFP^−^ cells (negative log2FC) (P<0.01, Table 1). The volcano plot shows the differentially expressed genes from the *Nr4a1*-EGFP mouse FACS sorted striatal cells (Fig. 1d). The *Nr4a1* log2FC indicates highly significant [p(adj) = 1.47E-113] enrichment in the striosome compartment but suggests by its lower value relative to other genes that the EGFP expression directed by the BAC transgene may not perfectly replicate endogenous *Nr4a1* expression.

Twenty-three differentially expressed genes were selected on the arbitrary threshold of log2 fold change greater than 7 and mean normalized counts greater than 40. These 23 genes together with *Nr4a1*, are shown in the heatmap (Fig. 1e). The heatmap includes *Irx1, Irx2* and *Foxf2* as top candidates for a role in defining the striosome.

To highlight transcription mechanisms responsible for defining the striosome MSNs, we identified the differentially expressed transcription factors using the mouse TF database (Table 1). The top 60 TFs enriched in each population according to their log2FC, i.e. EGFP^+^ population relative to the EGFP^−^ (upper panel), and in the EGFP^−^ population (lower panel) are shown in Fig. 1f. TFs enriched in the striosome include *Foxf2* as noted above and *Olig2* which are enriched 7 and 3-fold over their expression levels in the matrix, respectively. The gene sets that are enriched in the matrix compartment include numerous known and novel genes that may play a major role in the overall specification of the MSN and particularly matrix MSNs^14,18,31-40^. Known striatal TFs include *FoxP1, Dlx1,2,6, Gsx2, Mash1, Nkx2*.*1, Lhx6*, and *Lhx7*. The full list of TFs that define the striosome and matrix at this age are summarized in Table 1. 621 differentially expressed transcription factors were identified. Among these, 259 are enriched in the EGFP^+^ cells while 362 are enriched in the EGFP^−^ cells (presumed matrix cells).

As the *Irx1/2* genes were amongst the most enriched TFs in the striosomes, and have not previously been associated with the striatum, we carried out immunostaining of Irx1 to validate the predicted protein distribution. Irx1 localization corresponds to the DARPP-32 immunopositive striosomes in wild type PND3 mice, validating its predicted striosome enrichment (Fig. 1g). Further validation of a subset of the striosome factors and their function is described below. We did not characterize Irx2 immunohistochemistry as specific antibodies are not commercially available.

### Analysis of striosome enriched transcription networks defining *Foxf2* and *Olig2* as key differentiation factors

To identify potential master regulators that drive the transcription program of the striosome, we conducted transcription factor enrichment and co-expressor enrichment analysis with Enrichr^53,54^. We input the list of either differentially expressed genes or only differentially expressed transcription factors to obtain the significant upstream regulators and co-expressors for both gene sets (P<0.05). The intersect of the enrichment result and the differentially expressed transcription factors were further used as input to generate striosome and matrix specific transcription factor co-expression networks using GeneMANIA^55^ as a prediction tool (Fig. 2a,b). This allowed the generation of network reconstruction and expansion for striosome and matrix differentially expressed transcription factors, respectively. From the network analysis, we observed that *Nr4a1, Irx1, Olig2* and *Foxf2* interacted with a large number of transcription factors and these factors are enriched in the striosome (Fig. 2a). This indicates that these four TFs likely play a critical role in the development of striosomes and the MSNs located therein, as we have already shown for *Nr4a1*^22^. Since *Foxf2* and *Olig2* are potential novel regulators of the striosome (Fig. 1e,f), we evaluated whether their targets were coregulated in the striosome or the matrix. In the striosome, the transcriptional targets of *Foxf2* and *Olig2* are upregulated, whereas in the matrix, they are down-regulated (Fig. 2c,d). Thus, *Foxf2* and *Olig2* transcriptional targets are predicted to be enriched in the striosome. The TF network analysis of the matrix (Fig. 2b) indicates *Dlx1* and *Dlx2* are hub TFs (Fig. 2b). Potentially newly defined hub TFs in the matrix include *Pax7, Barhl2, Lhx9, Nhlh2, Eomes* and *Sp8* (Fig. 2b), the latter already associated with development of iMSNs^56^.

**Fig. 2:**
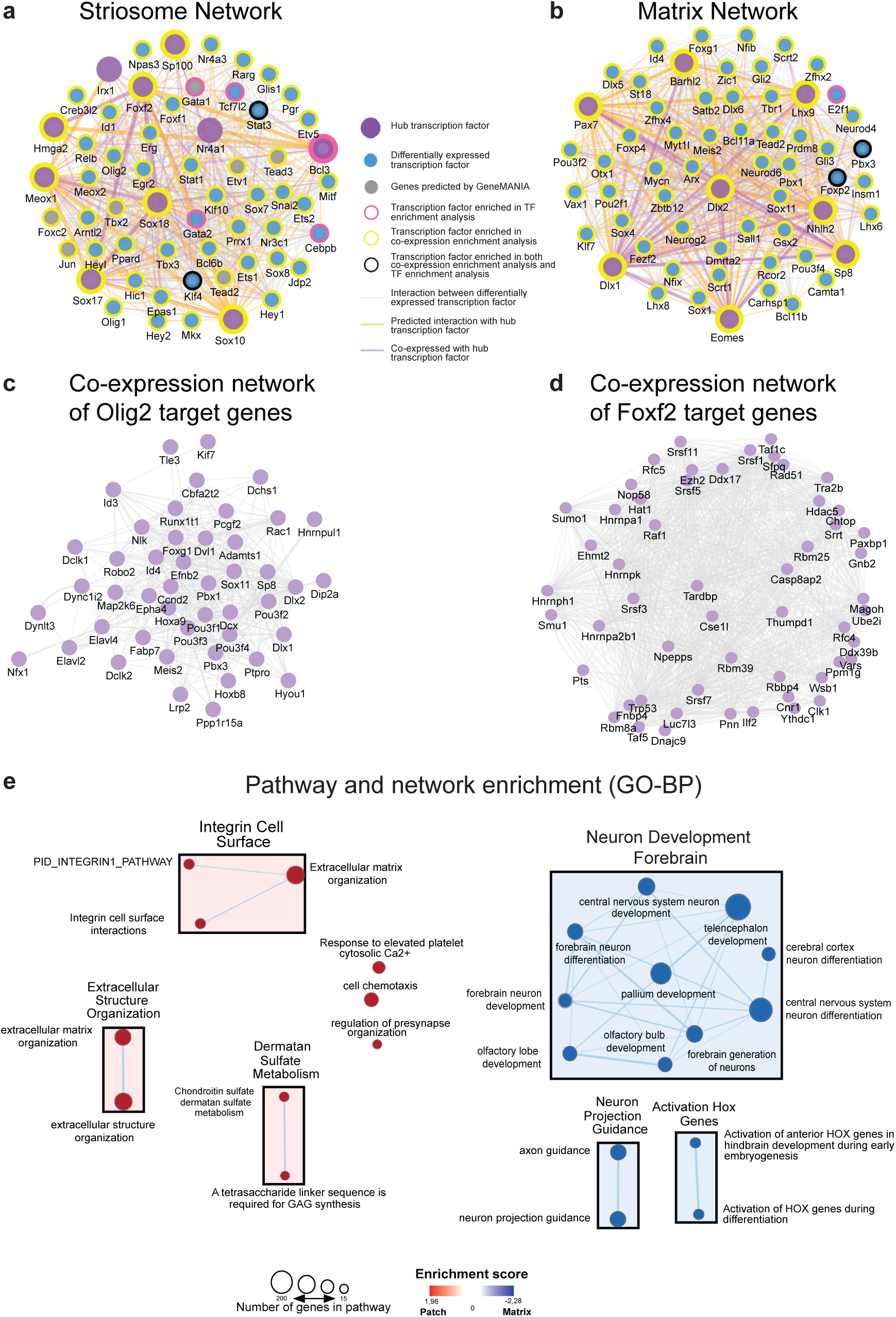
Transcriptomic networks and enrichment maps of striosome and matrix. **a**,**b)** Networks of transcription factors enriched in either striosome or matrix show interactions and co-expression among the regulatory genes. An arbitrary threshold of absolute log2 fold change greater than 5 was set to designate hub genes. **c, d)** Co-expression networks derived from striosome and matrix RNA-seq data show targets of Olig2 and Foxf2 are co-regulated in striosome and matrix. Genes with higher connectivity to other targeted genes are designated as hub genes. **e)** Pathway and network enrichment (GO-BP) for the striosome (left, red) and matrix (right, blue). The resulting enrichment map was annotated using the AutoAnnotate Cytoscape App.

In addition to defining the transcriptional network for the striosome and matrix, we analyzed the differentially expressed genes for overrepresented biological annotation or pathways, including GO term and KEGG pathway enrichment analysis^57^ (Table 1). Striosome enriched genes were associated with general development, including in the striatum, e.g. integrins, angiogenesis or cell motility, and nodes include “regulation of presynapse organization”. Matrix enriched genes were associated with neural system development and neurotransmission (Fig. 2e). Both *Nr4a1* and *Foxf2* are known to be associated with angiogenesis and/or establishment of the blood brain barrier^58,59^. Calcium regulation was associated with both gene sets. We also conducted a similar analysis with the differentially expressed transcription factor lists. For visualization and interaction network and GO enrichment analysis, we identified genes that are enriched in the striosome compartment, and most of these genes are involved in embryonic organ development (*Sox17, Gata3, Cebp, Nr4a3, Foxf1, Foxf2*), cell fate commitment (*SOX2, CEBP, Stat3, Olig1* and *Olig2*) and transcription activation such as RNA polymerase II binding transcription factors and enhancer binding (*Id1, Klf4, Tbx3, Erg2, Stat3*) indicating an active open chromatin structure (Table 1).

### ATAC-seq analysis defines compartment specific open chromatin regions in the striatum

To map chromatin accessibility of striosome and matrix cells, we combined fluorescence activated cell sorting (FACS) followed by preparation of 8 ATAC-seq libraries from 12 animals pooled in 4 independent samples. Overall, we obtained 322 million (average of 40.3 million) uniquely mapped reads after removing duplicate reads and those aligning to the mitochondrial genome (Supplementary Table 1). To quantitatively analyze differences among striosome and matrix cells, we generated a consensus set of 69,220 OCRs by taking the union of peaks called in the individual cells (**Methods**). We next quantified the number of reads which overlapped each OCR. Uniform Manifold Approximation and Projection for Dimension Reduction (UMAP) based clustering using the normalized read counts clearly separated striosome from matrix samples (Fig. 3a). Comparison of striosome vs. matrix peaks identified ∼44% of OCRs that were significant after multiple testing corrections (30,799 differentially modified OCRs at FDR ≤ 0.05) (Supplementary Table 2). Among these, 16,963 were striosome and 13,836 were matrix.

**Fig. 3:**
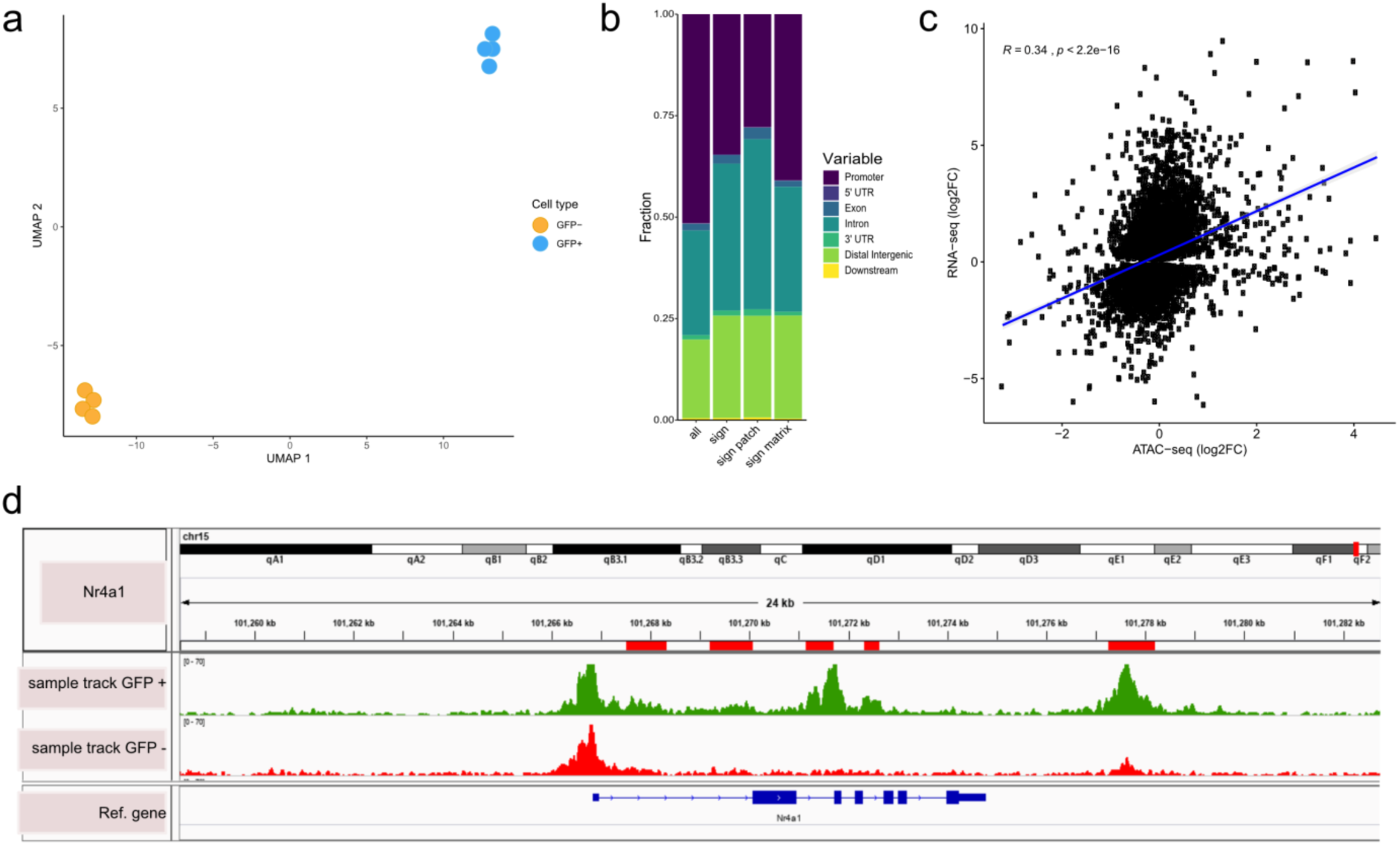
Open chromatin region (ATAC-seq) analysis identifies OCRs differentially located in striosome or matrix. **a)** Clustering of the individual samples using UMAP. **b)** Distribution of genomic features, i.e. location within gene (see Methods, Annotation of OCRs), of all and differential OCRs. **c)** Correlation of log2 fold changes in RNA-seq and ATAC-seq analyses. OCRs within TSS were considered for this analysis and only genes with adjusted P-value <0.01 from RNA-seq analysis is shown. **d)** *Nr4a1* ATAC-seq tracks in EGFP positive and negative populations showing the presence of a possible *Nr4a1* regulatory element highlighted by the red box in the EGFP^+^ sample.

We examined the location of OCRs with respect to the distance from transcription start site (TSS) and genic annotations. OCRs are in the vicinity of TSSs (Fig. 3b). The differentially accessible regions are enriched for non-promoter regulatory elements, suggesting a more important role for long-range regulation of gene expression in striosome and matrix specific OCRs. We tested the concordance of striosome and matrix specific genes and OCRs based on the RNA-seq and ATAC-seq analyses, respectively and found a high correlation (Pearson’s R = 0.34; P-value < 2.2×10^−16^) (Fig. 3c). In order to determine if the ATAC-seq could identify regulatory regions relevant to *Nr4a1*-EGFP enriched expression in the striosome, we compared the *Nr4a1* ATAC-seq tracks from PND3 *Nr4a1*-EGFP striatum FACs sorted in EGFP positive and negative populations for the presence of putative *Nr4a1* regulatory elements (Fig. 3d). This shows that a number of distinct OCRs within the *Nr4a1* gene are associated with the enriched EGFP^+^ cells. Finally, we used HOMER^60^ to identify which transcription factor binding motifs were selectively enriched in the OCRs of GFP^+^ and GFP^-^ populations. Within GFP^+^ OCRs, the top enriched motifs corresponded to the known consensus binding sequences for bZIP (Fra1/2, Fos, Atf3, JunB, BATF, AP-1, Fosl), HMG (Sox9, Sox10) and bHLH (Ap4, Tcf12, MyoD) family transcription factors. Notably, we identified *Olig2* motif among the top ranked enriched motifs, with a p-value of 1E-90 (Supplementary table 3).

**Table 3.**
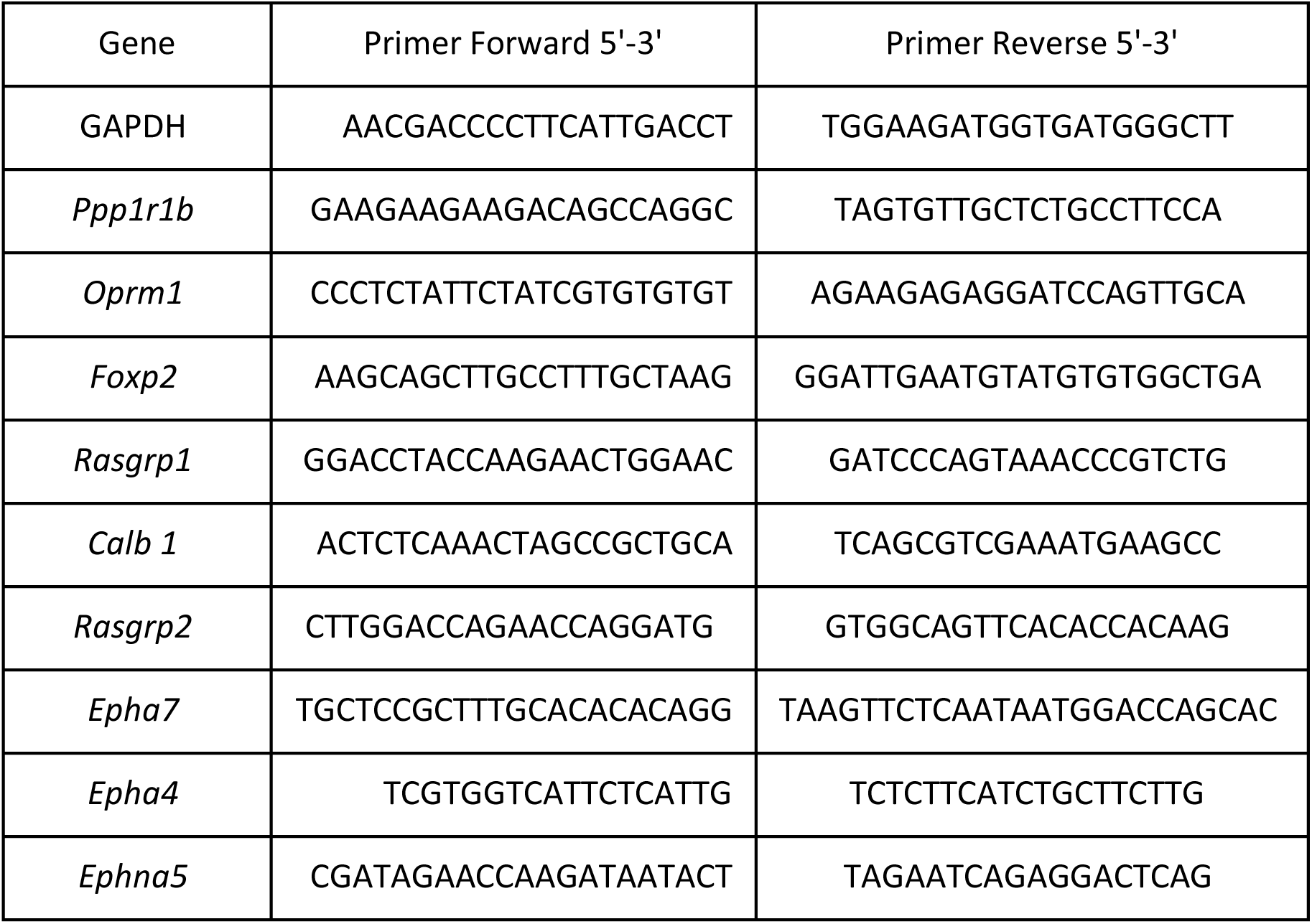
RT-qPCR murine primers sequence.

### *Foxf2* is required for striosome compartmentation

*Foxf2* is a member of the Fox protein family of TFs that regulate the expression of genes involved in embryonic development as well as adult life. It is expressed in brain endothelial cells and is critical for elements of craniofacial development, including cochlear development^61,62^. In our transcriptomic analysis, *Foxf2* is a hub gene (Fig. 2) and is ranked in the top twenty TFs associated with the striosome development at PND3. We confirmed with IHC that Foxf2 is colocalized with DARRP-32 striosomes on PND3 (Fig. 4a). Further, using RNAscope analysis of *Foxf2* mRNA, we also saw colocalization with *DARPP-32* mRNA (Fig. 4b). IHC and RNAscope failed to detect *Foxf2* expression in striatal neurons in the adult (Supplementary Fig. 1a). To determine whether *Foxf2* is required for early establishment of the striosome compartment, we analyzed wild type and *Foxf2*-null mice at E18.5 by IHC, as these mice are early postnatal lethal. Remarkably, the striosomes were absent in the *Foxf2* KO mice and thus *Foxf2* is necessary for the formation of the striosomes. DARPP-32 and TH immunostaining confirmed the lack of striosome assembly in *Foxf2* KO mice (Fig. 4c). It is important to note that the total expression level of DARPP-32 is not clearly decreased (Supplementary Fig. 1b), suggesting that the early born MSNs migrate to the striatal mantle, but other elements required for striosome compaction are impacted. In fact, the expression of members of Eph/Ephrin signaling system associated with control of brain cytoarchitecture ^63-67^ is reduced in *Foxf2* KO mice (Supplementary Fig. 1b).

**Fig. 4:**
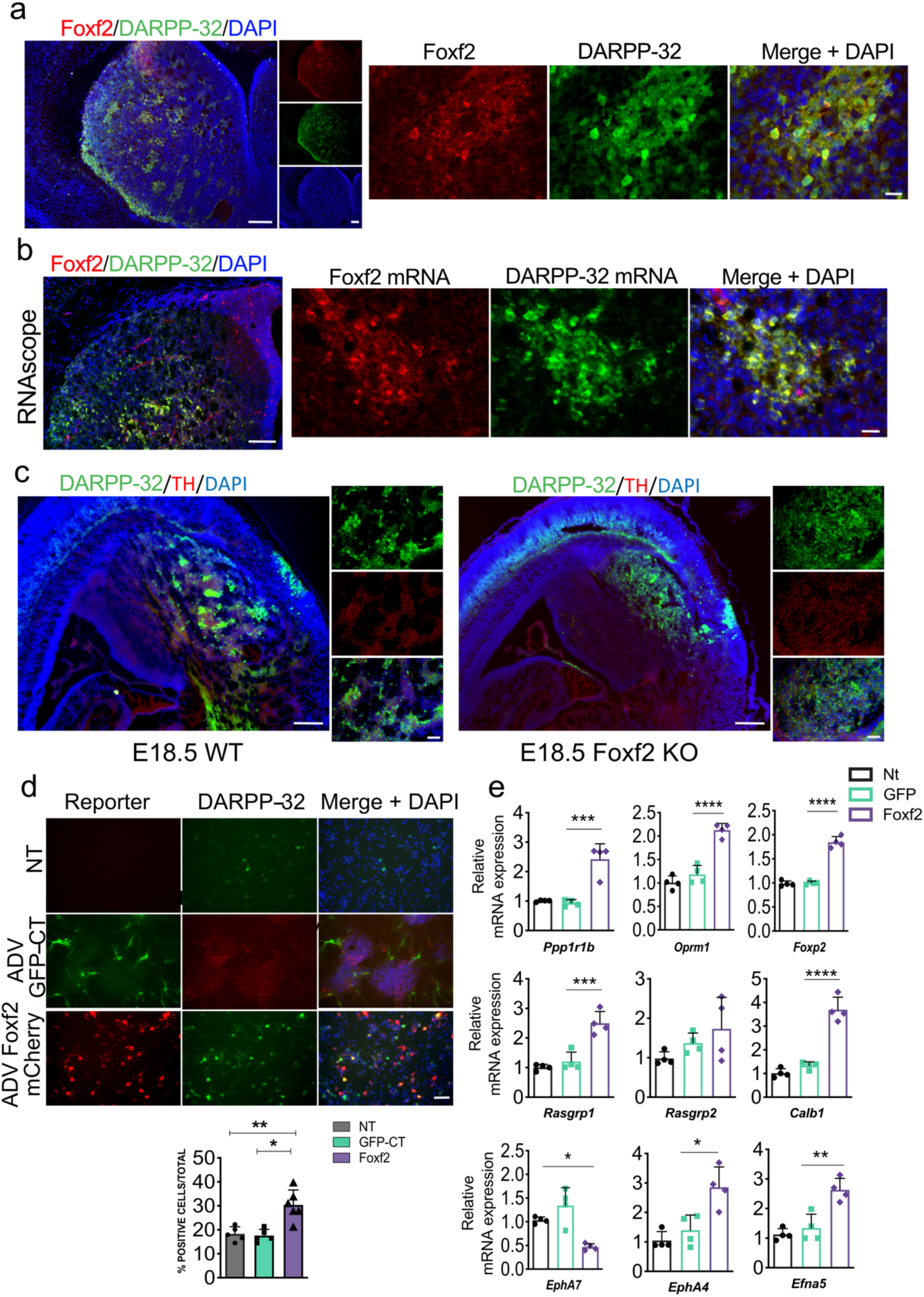
*Foxf2* is required for striosome compartmentation. **a)** Representative low and high-power micrographs of Foxf2 (red), DARPP-32 (green) and DAPI immunolabelling on 16 ⍰ ⍰ m thick coronal sections from wild type PND3 showing the localization of Foxf2 within the DARPP-32 positive striosomes. Scale bars correspond to 200 µm and 20 µm respectively. **b)** RNAscope analysis in 16 um thick coronal sections from wild type PND3 mice shows *Foxf2* mRNA co-localization with *Ppp1r1b*/DARPP-32 mRNA. Scale bars represent 100 µm and 20 µm. **c)** DARPP-32 (green) and tyrosine hydroxylase (red) immunostaining in sagittal sections of E18.5 WT and *Foxf2* KO mice show the absence of DARPP-32+, TH+ striosome assembly in *Foxf2* KO mice. Scale bars correspond to 100 µm and 50 µm respectively. **d)** Representative DARPP-32 staining in DIV9 WT primary striatal neurons transduced at DIV5 with adenovirus expressing GFP (control) or *FOXF2*-mCherry demonstrating the increase in DARPP-32 immunopositive cells in the cultures overexpressing Foxf2. Scale bar corresponds to 50 µm. **e)** Quantification of the percentage of DARPP-32 immunopositive cells in primary striatal neuronal cultures transduced with GFP or Foxf2 adenovirus. n=6 images from 3 individual cultures. One-way ANOVA corrected for multiple comparisons (Bonferroni’s) *P<0.05, **P<0.01. **f)** RT-qPCR assay shows increases in RNA of both striosome (*Ppp1r1b, Oprm1, Foxp2, Rasgrp1*) and matrix (*Calb1, Rasgrp2* and *EphA4*) markers in DIV-9 wild type primary striatal neurons 96h after transduction with ADV-Foxf2-mCherry. Importantly, FOXF2 overexpression decreases the mRNA of striosome ephrin receptor *Epha7* while increasing the levels of matrix *Epha4* and of its ligand *ephrin A5*. n=4 individual cultures, One-way ANOVA corrected for multiple comparisons (Bonferoni’s) *P<0.05, **P<0.01, ***P<0.001, ****P<0.0001. Error bars are standard deviation.

Given that *Foxf2* is required for the formation of the striosomes, we asked whether it is associated with the maturation of MSN gene expression. Overexpression of human FOXF2 via transduction with adenovirus promoted an increase in the percentage of DARPP-32 positive cells in primary mouse neuronal cultures derived from embryonic striatum (Fig. 4d). Further, analysis by RT-PCR demonstrated an increase in both striosome (*Ppp1r1b, Oprm1, Foxp2, Rasgrp1*) and matrix (*Calb1, Rasgrp2* and *EphA4*) marker mRNAs in wild type primary striatal neurons transduced with ADV-*FOXF2*-mCherry (Fig. 4e).

### *Olig2* is expressed in the striosome compartment and an associated OCR drives transgenic reporter expression in striosomes *in vivo*

Next, we evaluated the role of *Olig2* in the developing striosome compartment, including the validation of an associated striosome-specific OCR peak identified in our ATAC-seq analysis. IHC analysis of PND3 mice demonstrates that Olig2 co-localizes with DARPP-32 in (Fig. 5a). We confirmed this distribution using two antibodies raised against Olig2 including the one previously validated in *Olig2*-null mice (Supplementary Fig. 2a). Further, using RNAscope analysis of *Olig2* mRNA we found colocalization with *DARPP-32* mRNA (Fig. 5b,c). We did not evaluate the role of *Olig2* in the development of the striatum in *Olig2*-null mice due to early embryonic lethality. Note in the IHC that Olig2 protein is cytoplasmic in distribution in the striosomes, but is nuclear in scattered cells throughout the striatum, likely representing oligodendrocytes and their precursors.

**Fig. 5:**
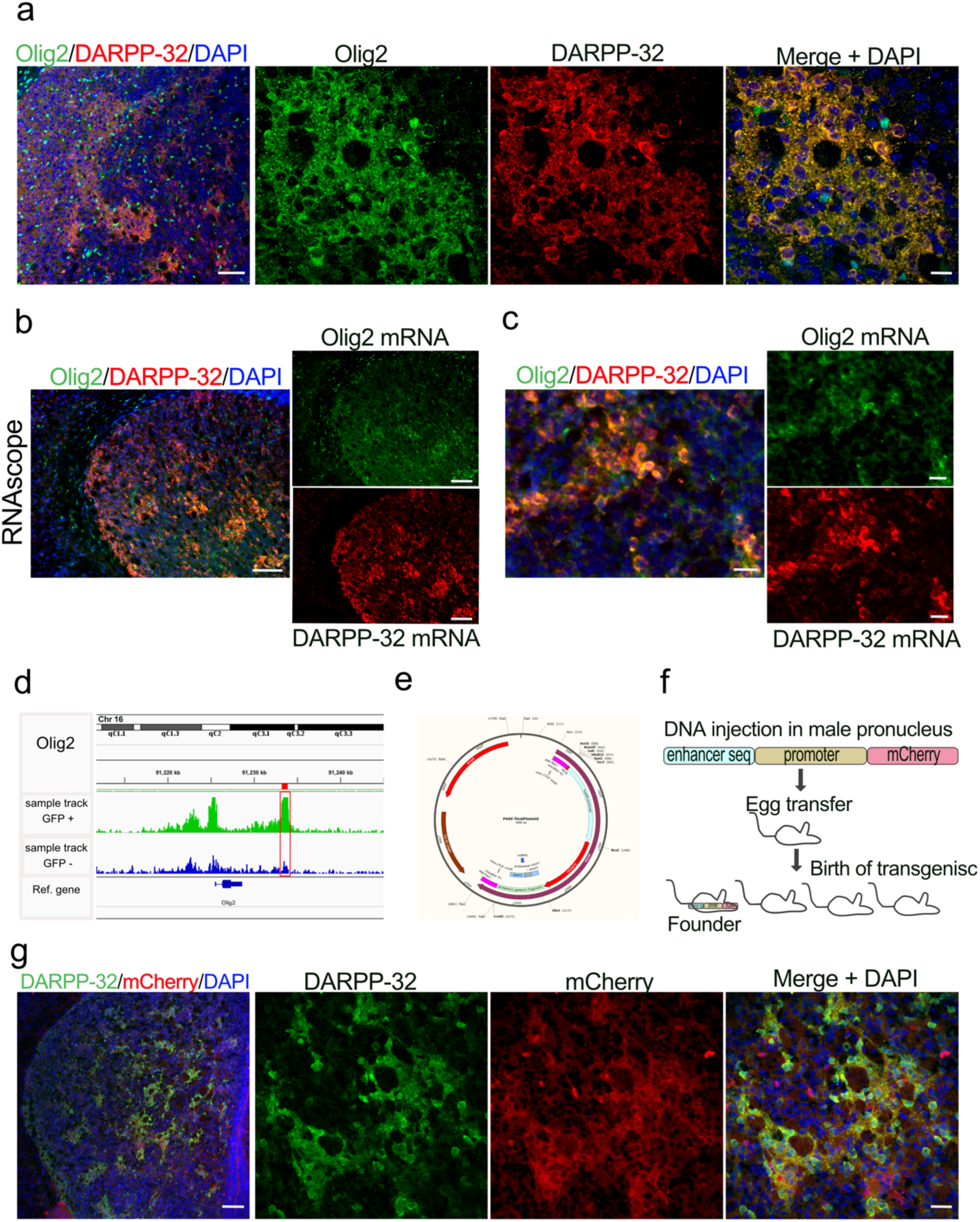
Olig2 is expressed in striosomes. and a downstream intergenic OCR drives transgenic reporter expression in striosomes. **a)** Representative low and high-power micrographs of Olig2 (green), DARPP-32 (red) and DAPI immunolabelling on WT PND3 16 ⍰m thick coronal sections showing the localization of Olig2 within the DARPP-32 positive striosomes. Scale bars correspond to 100 µm and 20 µm respectively. **b)** Low and **c)** high power magnification of RNAscope assay visualization of *Olig2* and *Ppp1r1b*/DARPP-32 mRNA indicating their colocalization in PND3 wild type striosomes. **d)** *Olig2* ATAC-seq tracks in EGFP^+^ and EGFP^−^ samples from PND3 *Nr4a1*-EGFP striatum showing the presence of an OCR downstream of the *Olig2* gene, restricted to EGFP^+^ samples (highlighted by the red box in the EGFP^+^ sample). **e)** Map of the p688/mcherry plasmid used for the cloning of the *Olig2* OCR shown in (d) for the generation of transgenics. **f)** Steps taken for the generation of transgenics expressing Olig2 enhancer: the insert containing the enhancer, the promoter and the reporter sequences is injected in male pronucleus, the eggs are transferred in a pseudo pregnant female that will give birth to potential founders containing the insert. **g)** Representative low and high-power micrographs of mCherry (red), DARPP-32 (green) and DAPI immunolabelling on 16 um thick coronal sections from PND3 *Olig2* -OCR transgenics indicating that the gene fragment acts as an enhancer and drives expression of the mCherry reporter specifically in striosomes. Scale bars correspond to 200 µm and 20 µm respectively.

We also wanted to determine if the ATAC-seq was able to identify functional, compartment-specific OCRs, outside promoter regions. We therefore sought to highlight a putative enhancer peak in a gene preferentially expressed in the striosomes. We compared the *Olig2* ATAC-seq tracks from EGFP^+^ and EGFP^−^ cells from FACs sorted PND3 *Nr4a1*-EGFP striatum and identified a peak several Kb downstream of the *Olig2* gene (Fig. 5d). We cloned the putative *Olig2* enhancer (chr16:91,233,041-91,234,111) into a p688 mCherry reporter plasmid and obtained 14 positive founders after pronuclear injection (Fig. 5e,f). mCherry was expressed in 9/9 founders on PND3 and co-localized with DARPP-32 (representative section Fig. 5g). The transgenic reporter was expressed in 2/5 founders assayed as adults but its expression was low and restricted to oligodendrocytes as indicated by the lack of co-localization with NeuN (Supplementary Fig. 2b, c). This analysis of an *Olig2* OCR demonstrates that the validated enhancer sequences from this rich data set will be useful in the development of tools to define and manipulate the striosome and matrix striatal compartments.

Next, we determined whether expression of OLIG2 in mouse primary MSNs promotes maturation towards striosome MSN fate (Fig. 6a). Expression of OLIG2 from transduced adenovirus does not increase the number of DARPP-32 immunopositive cells (Fig. 6b,c); however, it increased the mRNA for striosome markers *Oprm1, Foxp2* and *Rasgrp1* (Fig. 6d), but not the matrix marker, *Calb1*.

**Fig. 6:**
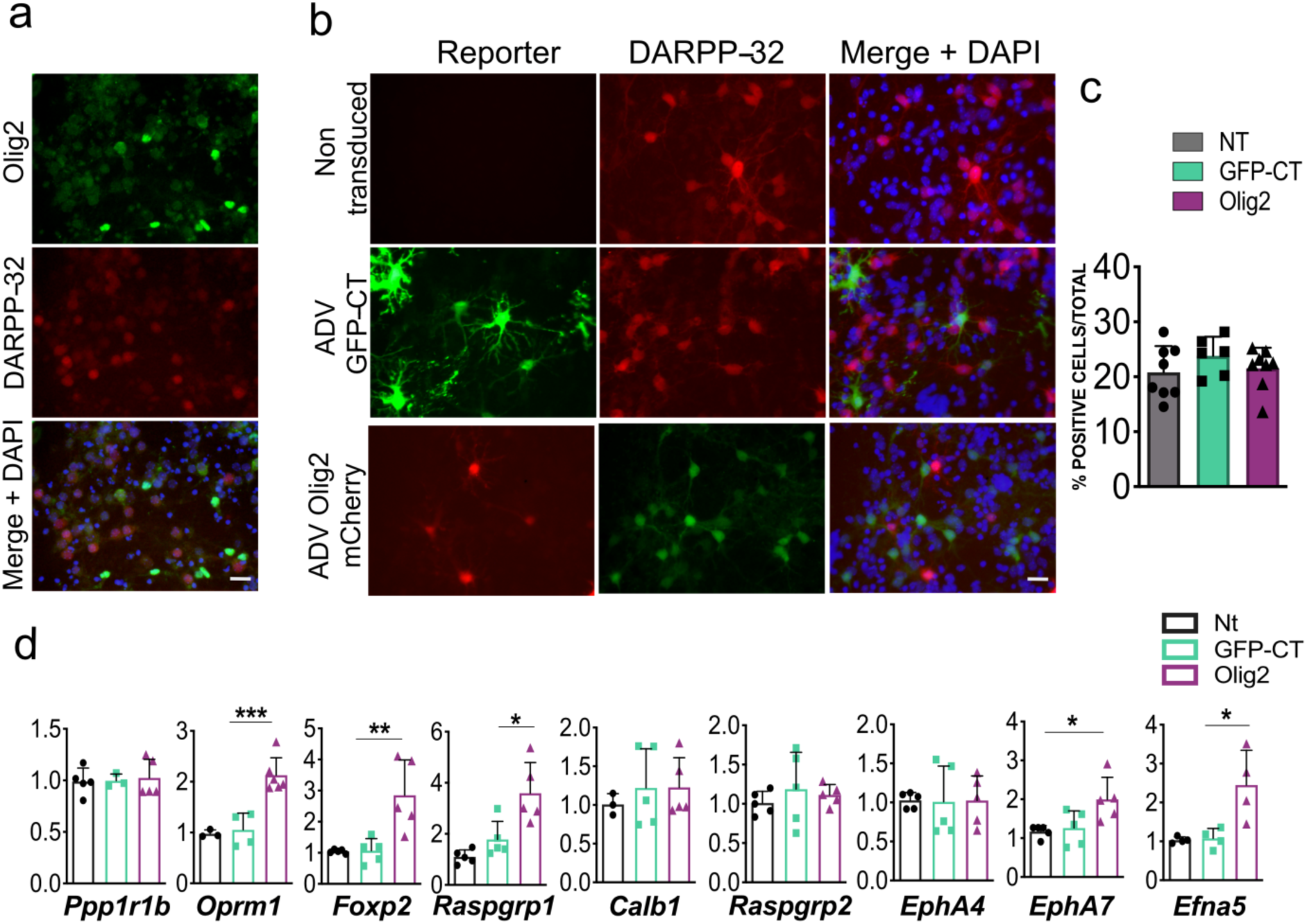
Olig2 is expressed in MSNs *in vitro* and its overexpression promotes their maturation towards a striosome phenotype. **a)** Representative staining in DIV9 WT primary striatal neurons showing the expression of Olig2 in DARPP-32 immunopositive MSNs. **b**,**c)** The number of DARPP-32 immunolabeledDIV9 WT primary striatal neurons is not increased following transduction for 96 h with ADV-*OLIG2* mCherry. ADV-GFP transduced and non-transduced cells were used as controls. Scale bar correspond to 50 µm. n=8 images from 3 individual cultures, One-way ANOVA, p > 0.05, non-significant. **d)** RT-qPCR assay shows increases of mRNA for striosome markers *Oprm1, Foxp2* and *Rasgrp1* in DIV9 wild type primary striatal neurons 96h after transduction with ADV-Olig2-mCherry. n=5 individual cultures, One-way ANOVA corrected for multiple comparisons (Bonferroni’s) *P<0.05, **P<0.01, ***P<0.001, ****P<0.0001. Error bars are standard deviation.

### Transcription network analysis defines STAT pathway as a key regulator

In order to quantitatively measure the importance of individual TFs in the striosome network (Fig. 2a), we used the arithmetic mean of the rank of each of the TFs at each node. The ranking was based on the number of edges expanding from each node and the number of overlapping TFs shared between nodes that interact with *Nr4a1*. Among the 58 TFs in the striosome network, the average rank of *Stat3* is 4.5, which ranks at the top. We also built the gene regulatory network with the entire differentially expressed TF set through the regulatory data curated by ORegAnno database^68^. This analysis resulted in the identification of *Stat1* as the TF that regulated the greatest number of differentially expressed TFs (314 out of 811) in our network (Fig. 7a). Thus, we identified that *Stat3* and *Stat1* appear to play an important role in the TF co-expression network and the gene regulatory network respectively, indicating that the STAT pathway might be a key regulating pathway during the differentiation of striosome neurons. Both *Stat1* and *Stat3* are enriched in the EGFP^+^ cells (Table 1). To experimentally test our bioinformatic analysis we expressed human STAT1 in primary mouse MSNs. STAT1 overexpression increases the number of DARPP-32 immunopositive cells (Fig. 7b,c) and increases the mRNA for both striosomal *Ppp1r1b, Oprm1, Foxp2, Foxf2* and *Olig2* as well as for the matrix marker *Calb1* (Fig. 7d). Particularly notable is the upregulation of other TFs identified in our analysis, i.e. *Foxf2* and *Olig2*.

**Fig. 7:**
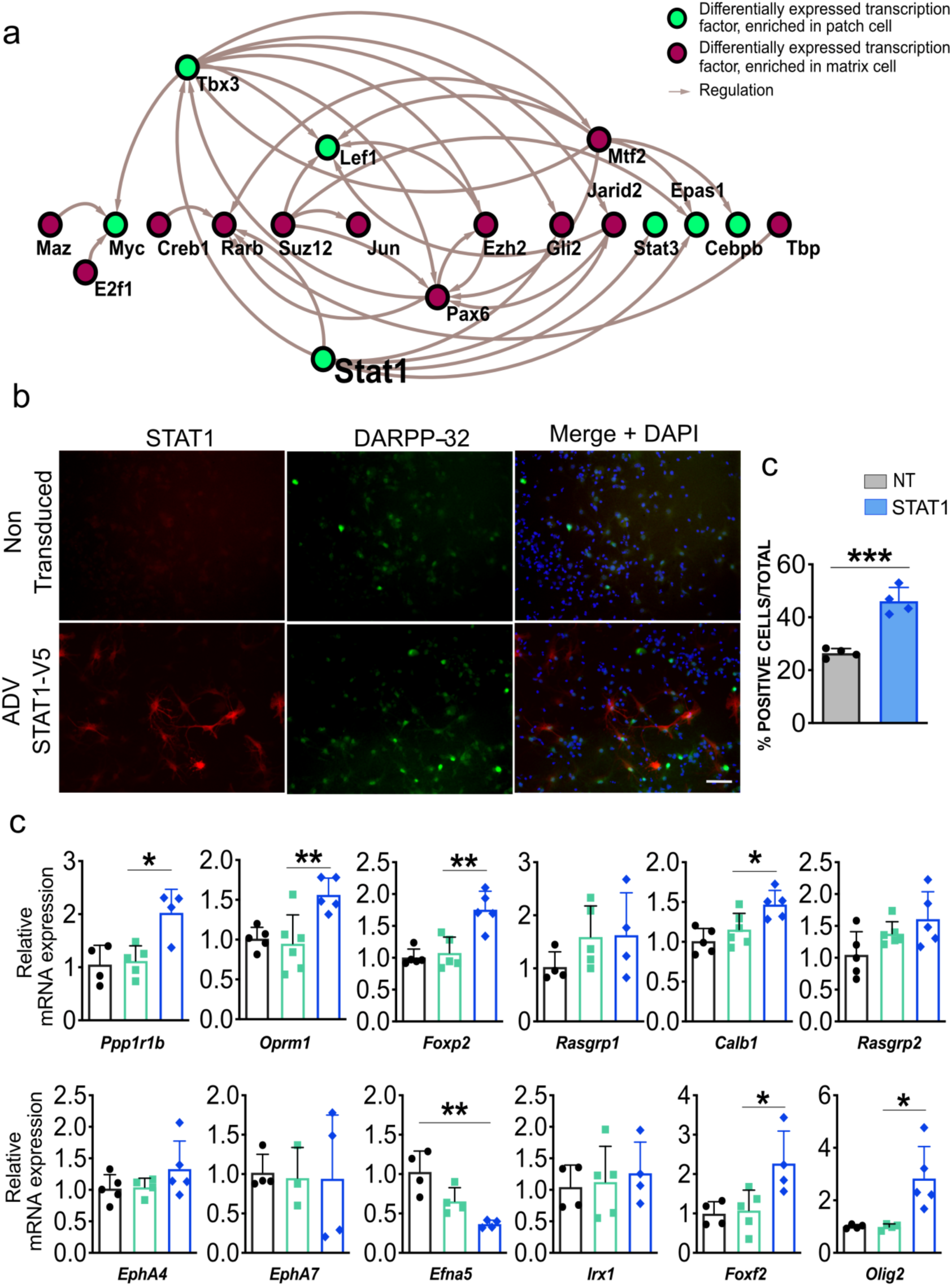
STAT1 overexpression in MSNs *in vitro* promotes their maturation and increases the levels of *Foxf2* and *Olig2* mRNA. **a)** Network demonstrating that *Stat1* may be a “master” TF in striosomes as it regulates the greatest number of TFs enriched in that compartment. **b**,**c)** Stat1 and DARPP-32 immunolabeling in DIV9 WT primary striatal neurons transduced for 96 h with ADV-*STAT1*-V5 vs. non-transduced controls show that Stat1 overexpression increases the number of DARPP-32 immunopositive cells. Scale bars correspond to 50 µm. n=4 images from 4 individual cultures, t-test ***P<0.001. **d)** RT-qPCR assay shows increases of mRNA for both striosome markers *Ppp1r1b, Oprm1, Foxp2, Foxf2* and *Olig2* as well as matrix marker*Calb1* in DIV9 wild type primary striatal neurons 96h after transduction with ADV-*STAT1*-V5. n=4 individual cultures, One-way ANOVA corrected for multiple comparisons (Bonferroni’s) *P<0.05, **P<0.01, ***P<0.001, ****P<0.0001. Error bars are standard deviation.

### Transcriptional regulation of striosome and matrix specific transcription networks is dysregulated in neurological diseases

HD is a monogenic movement disorder, has extensive transcriptomic data sets and has been linked to developmental dysregulation in the striatum^69-71^. Therefore, we evaluated if the genes expressed in striosomes or matrix in our data sets overlapped with HD transcriptomic public datasets. We applied the cell similarity test using the GEO database to obtain the differentially expressed published gene lists. We obtained twelve differentially expressed gene lists, and five of them were from human and seven of them were from mouse (Table 2, twelve RNA-Seq experiments). In addition to these twelve differentially expressed gene lists, we chose two mouse striatal microarray datasets and one dataset from human substantia nigra cells from a study focusing on Lewy Body Dementia, for a total of fifteen datasets for meta-analysis (Table 2).

In order to accurately compare the similarity between the different experiments, we considered the significance of the changes in expression levels of the gene and the expression pattern of the gene. We performed two computational experiments. First, we compared the experiments by only including genes that share the same expression pattern with striosome cells (Fisher’s Exact Test). Second, we compared only the genes that share the same expression pattern with matrix cells (Fisher’s Exact Test). Among the nine differentially expressed gene lists derived from mouse, six of them were significantly similar to striosome neurons and eight were significantly similar to matrix neurons (Table 2). Among these nine datasets, seven of them were significantly more similar to matrix cells than to striosome cells. Further, the two mouse datasets which showed weak similarity to matrix cells also showed weak similarity to striosome cells. From this analysis we can conclude that the transcriptomic correlation with mouse HD cells are more similar to the transcriptomic profile of matrix cells in mice. Among the six differentially expressed gene lists derived from human, four of them were significantly similar to striosome neurons and four of them were significantly similar to matrix. The motor cortex cell dataset showed weak similarity to both matrix and striosome cells. For the other five datasets, three of them showed stronger similarity to striosome cells than to matrix in the human transcriptomic data sets. Overall the dysregulated genes in mouse and human HD striatum overlap with our striosome and matrix transcriptomic data sets.

### TFs drive MSN fate iPSCs-derived NSCs

There is a critical need to generate MSN subtypes for disease modeling and there are no existing protocols that generate striosomal MSNs or use of TFs that specifically drive a striosome phenotype. To determine whether the FOXF2, OLIG2 and STAT pathways are critical for human development of MSNs and represent useful TFs for disease modeling of MSN subtypes, we expressed the TFs alone or in combination in NSCs derived from HD patient induced pluripotent stem cells (Fig. 8). STAT1 overexpression, with or without FOXF2, increases the number of OPRM1 immunopositive cells (Fig. 8a), with a trend in the same direction with OLIG2. STAT1 and OLIG2 together also increase the expression of PPP1R1B. Alone, only OLIG2 increases *RASGRP1* and *EPHA4*, mRNAs but in combination, STAT1 and FOXF2 are able to induce *<u>RASGRP1</u>*, although to a lesser extent. Interestingly, *RASGRP2*, a marker of the matrix, behaves inversely to that of *RASGRP1* to some extent, consistent with their expression *in vivo*. OLIG2 alone also increases the matrix marker *CALB1* mRNA, but at a lower-fold than the striosome markers. Notably, in contrast to its effect in primary MSNs, FOXF2 alone increased only *RASGRP2*, and not even the marker of pan-neuronal differentiation *TUJ1*, which along with the *in vivo* absence of striosome compartmentation in the striatum despite the presence of PPP1R1B neurons, may indicate an early role restricted to compartmentation and not overall phenotypic maturation as defined by expression of gene markers. Consistent with this hypothesis, FOXF2 alone failed to induce expression of *BCL11B* mRNA, a transcription factor required for striatal development ^31^ and decreased in HD^71,72^, or even *TUJ1*. Similar results were observed in isogenic controls C116-NSCs (Supplementary Fig. 3), in which OLIG2/STAT1 clearly induce OPRM1 expression. OLIG2 alone followed by OLIG2/STAT1 in combination may thus represent a treatment for induction of a striosome fate. It remains to be determined whether FOXF2 can actually induce compartmentation in 3D cultures.

**Fig. 8:**
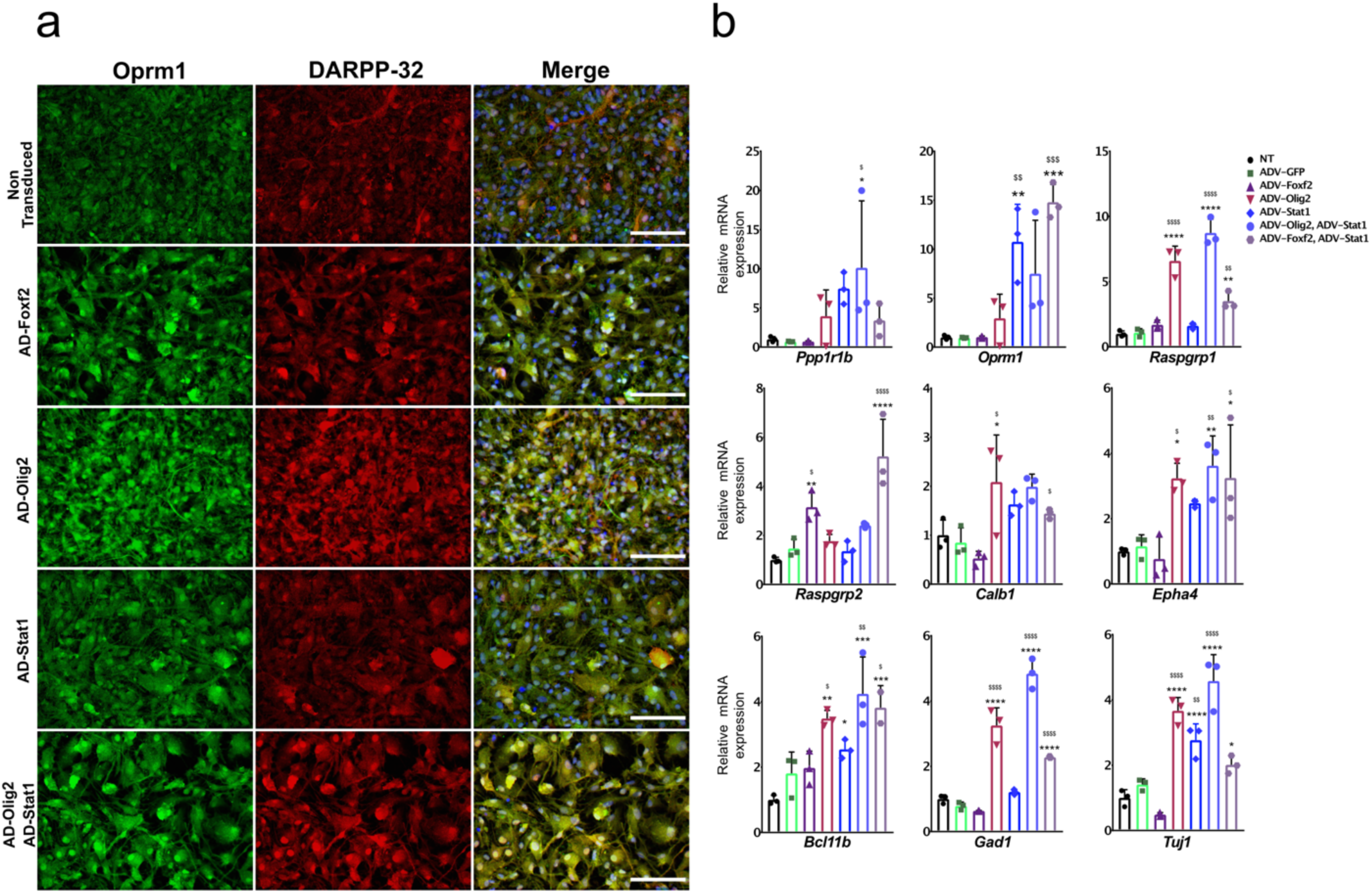
Foxf2, Olig2, and Stat1, alone and in combination, promote MSN differentiation in NSCs from human HD induced pluripotent stem cells. **a)** HD72-NSCs transduced for 4 days with ADV-*FOXF2*, ADV-*OLIG2* and ADV-*STAT1* were immunostained with Oprm1 (green) and DARPP-32 (red). Non-transduced cells and AD-GFP were used as control. Scale bars: 100 µm. **b)** RT-qPCR assay on HD72-NSCs transduced for 4 days with ADV-GFP, ADV-*FOXF2*, ADV-*OLIG2* and ADV-*STAT1*. n=3 individual cultures. One outlier (one of the three technical data points) was excluded in *Bcl11b* RT-qPCR when transduced with ADV-*FOXF2* and ADV*-STAT1*. One-way ANOVA for multiple comparisons (Dunnett’s) *P<0.05, **P<0.01, ***P<0.001, ****P<0.0001. * for comparison to non-transduced and $ for comparison to ADV-GFP. Error bars are standard deviation.

## Discussion

The mouse caudate nucleus develops from two periods of continuous neurogenesis and migration during the embryonic and early postnatal period^73,74^. During the first postnatal week, the striosomes are already compartmentalized, and were originally described as “dopamine islands” as determined by TH terminals^75^. Therefore, in the early part of the first postnatal week, maturation of the striosomes is advanced relative to the matrix, although this distinction has only been assayed for a handful of markers. We therefore hypothesized that selective sorting of *Nr4a1*-EGFP^+^ labelled striosome neurons at PND3 would lead to two main classes of cells, the first being young, but relatively mature, striosome neurons, and the second being relatively immature EGFP-negative neurons already committed to become matrix neurons. Our goals were to create RNA-seq and ATAC-seq databases which could provide the foundation for identification of further markers, including TFs, in early striosomes and developing matrix, and for derivation of regulatory networks for these two compartments. In addition, we aimed to identify TFs that could be used to better model striatal compartmentation in hiPSC systems and to develop tools to direct expression to neuronal subsets for their eventual *in vivo* manipulation and study.

For the first validation, we deliberately chose a TF highly enriched in the EGFP^+^ compartment, i.e. *Foxf2* (log2FC = 7.9), which has not previously been associated with neuronal development. In the *Foxf2*-null mouse, we demonstrate that the striatum shows a lack of normal striosome compartmentation in the presence of more diffuse expression of DARPP-32. Moreover, Foxf2 regulates the expression of Ephrin A5 and to some extent EphA7, both known to guide neuronal migration and connectivity of different areas of the brain^63,65,66,76-78^. In the context of gut development, components of extracellular matrix are severely reduced in the absence of *Foxf2*, leading to and lack of epithelial cell polarization and tissue assembly^79^. *Foxf2* is also required for correct establishment of cochlear cytoarchitecture^80^ controlling the arrangement of type II spiral ganglion neuron innervation and the density of axons throughout the organ of Corti. *Foxf2* constitutive deletion also clearly has wide-ranging effects on brain development beyond the striatum, a subject for further investigation. *In vitro* in primary MSNs, increased *Foxf2* induces genes expressed in striosomes and matrix but not in the less mature hNSCs, which together with the *in vivo* appearance of the *Foxf2*-null striatum, suggests that in the embryonic period, *Foxf2* alters genes associated with sorting of striosomes and matrix neurons, and not necessarily with MSN-specific maturation. The precise molecules and mechanisms governing striosome/matrix development and connectivity remain a subject of research^44,81,82^, but the notable effect of *Foxf2* overexpression on components of the ephrin system imply regulation of signals required for compartmentation^64,78^.

Amongst the TFs enriched in the EGFP^+^ compartment, there were activators and others that are more commonly described as transcriptional repressors, e.g. *Zfp48*8. This zinc finger protein is of particular interest as it cooperates with Olig2^83^. *Olig2* (log2FC = 3.05) was not amongst the most enriched in its group and the gene family is of course most associated with OL lineage. In fact, however, both *Olig1* and *Olig2* are also regulators of neuronal subtype development in the human fetal central nervous system^84^ and in mouse, *Olig2* is regulated by *Dlx1* and *Dlx2* and appears to be involved in the neuron-glial switch^85^. Specifically, amongst other roles, *Olig2* participates in the development of GABAergic neurons in the pre-thalamus and thalamus^86-88^. Although we are unable to assay development of the striatum in *Olig2*-null mice due to early lethality, and therefore are unable to state that *Olig2* is required for striosome development, our data demonstrate the expression of *Olig2* in early striosomes, and its ability *in vitro* to induce a related phenotype. Notably, Olig2 localized to the cytoplasm in striosome neurons, where it has been shown to act to regulate differentiation of astrocytes^89^. The validation of the predicted OCR in the *Olig2* gene as a striosome-specific enhancer in the striatum lends further support to the function of *Olig2* and also highlights the value of the ATAC-seq database.

Based on the known expression pattern of *Nr4a1*, we were not anticipating that either set of cells would be purely neuronal. *Nr4a1* is expressed in endothelial cells^90^, and in cells of the myeloid lineage^91^. Published transcriptomic studies reveal expression of *Nr4a1* in OPCs which is downregulated with maturation^92^. *Nr4a1* is also expressed in astrocytes, particularly when activated as an immediate early gene^93^.

The GO analysis of the EGFP^+^ positive compartment highlights enrichment of genes and pathways associated with vasculature development, regulation of angiogenesis and extracellular structure organization. Notably, it has been suggested that the vasculature may play a role in striatal compartmentation, as striosomes appear to have higher vascularization compared to matrix ^94^. The importance of the blood vessels in regulation of cortical neuronal migration and maturation is well described^95, 96, 97^ and vascular and neuronal development share common signaling pathways^98,99, 100^ including netrins, semaphorines and members of vascular endothelial growth factor (VEGF) family.

The use of these new data bases to construct a transcriptional regulatory network also led to the identification of *Stat1/3* as a TF able to modulate striosome and matrix phenotypes. Notably, these two members of the Stat gene family can dimerize with each other^101^. Neither Stat1 or 3 has been previously associated with MSN development, but they are reported to promote neuronal differentiation^102^. Further work is required to determine the role of *Stat1* in striatal development *in vivo*, but we did find a clear maturation effect *in vitro*. The effects of validated TFs on members of the ephrin family highlights the fact that although we focused on TFs, these data sets also identify compartment-enriched encoding proteins other than TFs which are likely critical for establishing and maintaining striatal compartmentation.

The transcription regulatory networks generated for each cell compartment have defined other potential mechanisms regulating striosome and matrix cell fate. The co-expression enrichment analysis has highlighted a set of TFs that coexist with *Foxf2* and may play a role in striatal MSN differentiation, including *Sox7, Sox17, Sox18, PrrX1, Tead2, Bcl6, Klf4, Egr2*, and *Tbx2*. Other than Tead2 and Tbx2, these TFs are enriched in the EGFP^+^ cells. *Sox17* belongs to the *Sox* family of TFs known to regulate oligodendrocyte progenitor cell expansion and differentiation during development and repair via Notch signaling^103^. Both *Sox17* and *Notch* signaling regulate *TCF7L2* expression^103^, which is enriched in the same compartment. *TCF7L2* regulates calcium signaling^104^, a function associated with both gene sets. In spinal cord patterning, *TCF7L2* in partnership with *Tcf4* can repress *Olig2* expression through recruitment of HDAC activity and is required for cell fate specification^105^. Histone deacetylase activity likewise plays a role in MSN maturation^106^. For *Olig2*, the TF co-expression network highlights *Etv5, Mitf, Egr2, Sox10, Olig1, Hey1*, and *Arntl2*, which are also mostly enriched in the EGFP^+^ compartment. There is considerable evidence of cooperation between *Olig2* and *Olig1* in other systems, including neuronal^107^. Thus, detailed analyses of these databases suggest multiple TF networks involved in striosome development which warrant further investigation. The same is likely true for the matrix compartment.

The enrichment of OPC markers in the EGFP^+^ compartment was particularly prominent, and these genes included TFs, structural genes, e.g. *Gpr17*, and metabolic genes, e.g. *Enpp7*. Although beyond the scope of this report, the role of *Nr4a1* in oligodendroglial development should be further investigated, and based on the Olig2 data, the possible role of some of these genes in the neuronal lineage will also be of interest, as highlighted in the preceding paragraph.

Consistent with our hypothesis, the EGFP^−^ cells were indeed enriched in markers of immature MSNs. Most notably, these included multiple members of the *Dlx* family, required for patterning and differentiation^34^, *Isl1* which promotes commitment of striatonigral neurons^108^, and markers of neuronal differentiation but not specifically related to the striatum, e.g. *Sox11* and *Pax6*, which is expressed in the ventricular zone of the lateral ganglionic eminence^109^. The *Arx* family is involved in neuronal progenitor proliferation and was identified by its association with lissencephaly syndromes, some of which have specific basal ganglia deficits^110^. *Eomes/Tbr2* is associated with cortical development, as is *Tbr1*, raising the possibility of contamination by cortical elements during dissection and cell selection^109^. It should be noted, however, that both *Tbr1* and *Satb2* mRNAs are detected in the striatum [Allen Mouse Brain Atlas (mouse.brain-map.org)]. Interestingly, *Foxp2* and *Oprm1* are associated with striosomes, yet are enriched in the EGFP^−^ cell compartment^4^, while *Ppp1r1b*, a commonly used early marker of striosomes, is only slightly enriched in the EGFP^+^ compartment. These data imply that in some cases, there may be a dissociation between transcription and translation at this stage, although it is known that *Oprm1* mRNA distribution is initially diffuse during striatal development^111^. Other factors enriched in the EGFP^−^ compartment have not previously been associated with neuronal differentiation, or other cell types in the CNS, e.g. *Fank1*, or have been found in the brain but not in the striatum, e.g. *Insm1*. We will continue to investigate such factors for their role in striatal development.

An important finding for our studies is the ability of some of these factors to drive the expression of MSN maturation in human NSCs derived from HD-iPSCs. We do not know if MSNs used for disease modeling have similar identities to those formed during human development, nor have we developed conditions to make pure subpopulations of MSNs (e.g., striatonigral, striatopallidal, striosome, and matrix subpopulations). Thus, we have established experimental and transcriptomic data sets that will allow us to improve existing protocols to further model striatal phenotype in human movement and cognitive disorders associated with MSNs in the developmental and adult brain.

An important question that remains is whether the TFs enriched in the EGFP^+^ compartment are terminal selectors, as defined by Hobert^46^ to include “coregulation of terminal effector batteries, combinatorial control mechanisms, and the coupling of initiation and maintenance of neuronal identity”. For the two TFs which we validated in detail, *Foxf2* and *Olig2*, their functions do not appear to include maintenance of striosomal identity, as their expression is undetectable in the adult mouse and there are no assigned striatal ATAC-seq peaks from human striatum^112^. Most TFs identified to date that are expressed both during MSN differentiation and in the adult, e.g. *Bcl11b, Foxp2*, and *Foxp1*, have not been studied specifically for gene expression alterations following their deletion in the adult, although it has been shown that siRNA knockdown of *Foxp2* in the adult striatum leads to motor and electrophysiological dysfunction^113^. Similar experiments to those we performed at PND3 will be performed in the adult to develop complementary data bases.

Together, we present two data bases derived from PND3 striatum comparing gene expression and open chromatin regions in developing striosome and matrix, demonstrating clear gene expression and epigenetic distinctions. Using these data sets, we describe a novel pathway via which Stat1 appears to regulate a transcriptional hierarchy which includes Foxf2 and Olig2, critical for striosome compartmentation and phenotypic maturation. Further, we demonstrate that these TFs can be utilized to better model striatal striosome MSNs in hiPSC systems and to develop genetic tools to direct expression to neuronal subsets for their eventual *in vivo* manipulation and study.

## Supporting information

Table 1 Differential expression Bioinformatics

## Methods

### Animals

Animal procedures were conducted in accordance with the NIH Guidelines for the Care and Use of Experimental Animals and were approved by the Institutional Animal Care and Use Committee of our institutions. The *Nr4a1-*EGFP mice used for this study were obtained from GENSAT. *Foxf2*^-/-^ and littermate wild-type (WT) controls at E18.5 were kindly provided by Dr. Peter Carlsson as well as and homozygote *Foxf2*^fl/fl1^. Mice were given *ad libitum* access to food and water and housed under a 12-h light/dark cycle. Both male and female mice were used in these studies.

### Enzymatic dissociation and FACS purification of striatal neurons

PND3 *Nr4a1*-EGFP heterozygous mice were obtained from a time set homozygote *Nr4a1*-EGFP bred to WT pairs. The pups were separated by sex, rapidly euthanized by decapitation, and the striata dissected under the microscope in ice-cold Hibernate-A medium (A1247501, Gibco, Thermo Fisher). The striata from 3 PND3 mice were collected per sample and exposed to enzymatic dissociation with papain as described in Lobo et al.^114^ with minor modification. Briefly, the tissue was incubated with frequent agitation at 37°C for 20 min in 2 ml of 2 mg/ml papain (Sigma, P4762) solution in Hibernate-A. The tissue was washed twice with 5 ml Hibernate-A including 1 mg/ml protease inhibitor (Thermo Fisher, 78442) and briefly triturated in 2 ml of Hibernate-A medium using fire-polished glass Pasteur pipettes until a single-cell suspension was obtained. The cell suspension was filtered through 70-µm mesh previously equilibrated with 2 ml of Hibernate-A. The cells left on the filter were collected by washing the filter with 1 ml of Hibernate-A. The cells were centrifuged for 5 min at 700g and treated for 10 min with DAPI (4’,6-diamidino-2-phenylindole) (1:4000, Sigma, 62248) to label dead cells. The cells were washed with 10 ml of Hibernate-A, centrifuged for 5 min at 700g and resuspended in 0.35 ml of Hibernate-A. Viable cells (DAPI negative) were sorted and collected into two populations: EGFP^+^ and EGFP^−^ using a BD Influx FACS sorter based on fluorescein-5-isothiocyanate (FITC) channel signal for EGFP. WT striatal cells were used to calibrate the FITC and DAPI signals.

The cells for RNA-seq (1000 cells/sample) were collected in 150 μl of Arcturus PicoPure extraction buffer. After the sorting, the samples were incubated at 42°C for 30 min and stored at -80°C until RNA extraction was performed using Arturus PicoPure RNA Isolation kit (Applied Biosystems, 12204-01). The cells for ATAC-seq (25,000 cells/sample) were collected in 1.5 ml Eppendorf tubes previously coated with 5% BSA, centrifuged at 300g for 10 min at 4°C and the pellet was then stored.

### Tissue preparation and immunofluorescence

PND3 mice were rapidly euthanized by decapitation, and brains were removed, washed in ice-cold PBS, and post-fixed for 24 h at 4°C in 4% PFA. The brains were then incubated in 30% sucrose/1X PBS for 24 h at 4°C and cryopreserved in OCT embedding medium (4583, Tissue-Tek Sakura). Serial coronal section (16 µm) were cut on a Leica cryostat, collected on Superfrost Plus microscope slides (Fisher Scientific) and frozen at - 20°C. Immunofluorescence was performed as previously described^3^. Sections were incubated with mouse anti-DARPP-32 (1:250, sc-271111, Santa Cruz Biotechnology), sheep anti-Foxf2 (1:2000, AF6988, R&D), rabbit anti-Olig2 (1:500, ab136253, Abcam), rabbit anti-Olig2 (1:8000, gift Dr. Wichterle)^115^, mouse anti-Olig2-Alexa488 (1:1000, MABN50A4, Millipore) rabbit anti-Irx1 (1:2000, PA5-36256, Thermo Fisher Scientific) or rabbit anti-Tyrosine hydroxylase (1:1000, OPA1-04050, Thermo Fisher Scientific) antibodies. The respective secondary antibodies included: anti-mouse Alexa 488 (1:400, A-11008, Thermo Fisher Scientific), anti-mouse Alexa 594 (1:400, A-11005, Thermo Fisher Scientific), anti-rabbit Alexa 488 (1:400, A-11034, Thermo Fisher Scientific), anti-rabbit Alexa 594 (1:400, A-11012, Thermo Fisher Scientific) or anti-sheep Alexa 594 (1:400, A-11016, Thermo Fisher Scientific). Sections were sealed with Vectashield hard-set mounting medium (H-1400, Vector Laboratories). Images were acquired using an Olympus BX61 microscope or a Confocal Zeiss LSM 510.

### Fluorescent in situ hybridization (FISH) using RNAscope technology

Postnatal day 3 WT brains were fixed in freshly prepared, ice-cold 4% PFA for 24 h at 4°C followed by equilibration in 10% sucrose gradient, then 20% and finally 30%, each time allowing the tissue to sink to the bottom of the container. The tissue was embedded in OCT compound (4583, Tissue-Tek Sakura) and stored at -80°C until sectioned. 16 µm-thick sections were cut using a Leica cryostat, collected onto Superfrost Plus slides maintained at -20°C during the sectioning. The slides were then stored at -80°C. RNAscope®Probe murine Mm-DARPP-32-C1 Ppp1r1b (NM_144828.1, bp590-1674, 405901), Mm-Olig2-C3 (NM_016967.2, bp865-2384, 447091-C3) and Mm-Foxf2-C2 (NM_010225.2, bp846-2316, 473391-C2)-were purchased from Advanced Cell Diagnostics probe catalog (ACD). For signal detection, we used Opal 520 and Opal 690 TSA plus fluorophores (Akoya Biosciences). The RNAscope Multiplex Fluorescent Reagent Kit v2 (ACD, 323100) used here provides the target retrieval solution, hydrogen peroxide, protease III, amplification reagents (Amp1-3), HRP reagents, DAPI, TSA buffer and wash buffer. We used a modified version of the manufacturer’s protocol for sample preparation, probe hybridization, and signal detection. Briefly, the fresh frozen sections on slides were retrieved from -80°C and briefly immersed in 1X PBS to wash off the O.C.T and then baked at 60°C for 30 min. Slides were then post-fixed in fresh 4% PFA for 1h at room temperature (RT). After fixation, the sections were dehydrated in a series of ethanol solutions (5 min each in 50%, 70%, and two changes of 100% ethanol) at RT and left to dry for 5 min at RT. Sections were treated with hydrogen peroxide for 10 min at RT and washed twice with distilled water. Subsequently, target retrieval was performed by boiling the slides for 5 min in 1X Target Retrieval Reagent (ACD), washed in distilled water, immersed in 100% ethanol and air-dried for 5 min at RT. A hydrophobic barrier was created around the section using an ImmEdge Pen (ACD, 310018) and completely dried at RT before proceeding to the next step. Sections were then treated with protease III for 5 min at 40°C in the pre-warmed ACD HybEZ II Hybridization System (ACD, 321721) inside the HybEZ Humidity Control Tray (ACD, 310012) and washed twice with distilled water. The *Foxf2*-C2 or *Olig2*-C3 probes were diluted at 1:50 in *DARPP32*-C1 probe. The sections were then hybridized with the probes, *DARPP-32* and *Olig2* or *DARPP-32* and *Foxf2*, at 40°C for 2 h in the HybEZ Oven (ACD), washed twice with 1X wash buffer, and stored overnight at RT in 5x SSC buffer (Thermo Fisher Scientific). The next day, the slides were rinsed twice with wash buffer for 2 min each, followed by the three amplification steps (AMP 1, AMP 2, and AMP 3 at 40°C for 30, 30, and 15 min respectively, with two washes with wash buffer after each amplification step). The signal was developed by treating the sections in sequence with the HRP reagent corresponding to each channel (e.g. HRP-C1) at 40°C for 15 min, followed by the TSA Plus fluorophore assigned to the probe channel (Opal 690 for DARP32-C1 probe at 1:2000 dilution and Opal 520 for *Foxf2*-C or *Olig2*-C3 probes, prepared at a dilution of 1:1500) at 40°C for 30 min, and HRP blocker at 40°C for 15 min, again with two wash steps after each of the incubation steps. Finally, the slides were counterstained with DAPI for 30 s, mounted using ProLong Gold mounting medium (Thermo Fisher Scientific) and stored at 4°C until ready for imaging. The sections were imaged using a 10x/0.3 N.A., 20x/0.8 N.A. or 40x/0.75 N.A. objectives on an AxioImager Z2 microscope (Carl Zeiss), equipped with a Zeiss Axiocam 503, and operated with Zeiss Zen Blue software (Carl Zeiss). Camera exposure times were set for all three channels (red for Opal 690, blue for DAPI and green for Opal 520) and were identical for all slides within each experiment. ImageJ was used for adjusting the brightness and contrast of the images.

### Primary neuronal cultures

E16.5 embryos were obtained from WT Swiss Webster timed bred females purchased from Charles River. E16.5 striatum was removed from by microdissection in cold Leibovitz’s medium (L-15) (Gibco-Invitrogen,11415064) and primary medium spiny neuronal cultures were prepared as described in^22^. Briefly, the tissue was incubated in Ca^2+^/Mg^2+^-free Hanks’ balanced salt solution (Sigma, 55021C) for 10 min at 37°C. The incubation mixture was replaced with 0.1 mg/ml papain in Hibernate E/Ca^2+^ (BrainBits) incubated for 8 min and rinsed in Dulbecco’s minimum essential medium (Gibco-Invitrogen, 21013024) with 20% fetal bovine serum (Gibco-Invitrogen, 10438026) and twice in Leibovitz’s medium (L-15). The tissue was then suspended in Dulbecco’s minimum essential medium with 10% fetal bovine serum, glucose (6 mg/mL) (Sigma, G7021), glutamine (1.4 mM) (Gibco-Invitrogen, 25030081) and penicillin/streptomycin (100 U/mL) (Gibco-Invitrogen, 15140122). Cells were triturated through a glass bore pipette and plated onto either Lab Tek eight-well slides (1.25 ×10^5^ cells/well) for immunocytochemistry or 24-well plates (1×10^6^ cells/well) for RT-PCR analysis, each previously coated with polymerized polyornithine (0.1 mg/mL in 15 mM borate buffer, pH 8.4) and air-dried. One h later, the media was replaced with Neurobasal (Gibco-Invitrogen, 21103049) supplemented with B27, (Gibco-Invitrogen, 17504044), GLUTAMAX (Gibco-Invitrogen, 35050061) and penicillin/streptomycin. The medium was changed every two days and the cells were assayed on day *in vitro* DIV 9.

### Neuronal adenovirus (ADV) transduction

ADV-CMV-*OLIGO2*-mCherry, ADV-CMV-*FOXF2*-mCherry, ADV-CMV-*STAT1*-V5 and ADV-CMV-EGFP were produced by SignaGen Laboratories using the human cDNA sequence. Viral transduction was performed at DIV5 with a multiplicity of infection (MOI) of 20. The virus was added in fresh medium, and the medium was changed 18 h later. Cells were harvested or fixed 96 h following addition of virus.

### Mouse neuronal immunocytochemistry

Cells were fixed in 4% paraformaldehyde in 0.1 M phosphate buffer, pH 7.4, and immunolabeled with mouse anti-DARPP-32 (1:250, sc-271111, Santa Cruz Biotechnology), rabbit anti-Olig2 (1:500, ab136253, Abcam) or rabbit anti-STAT1 (1:400, 14994S, Cell Signaling Technology) followed by anti-mouse Alexa 488 (1:400, A-11008, Thermo Fisher Scientific) and anti-rabbit Alexa 594 (1:400, A-11012, Thermo Fisher Scientific). To identify the total number of cells the nuclei were stained with DAPI (4’, 6-Diamino-2-phenylindole dihydrochloride) (1:10000, Millipore-Sigma). Images were acquired using Olympus BX61 microscope and analyzed using Fiji software (ImageJ).

### Real-time qPCR for mouse cells

RNA from DIV9 primary MSNs was extracted with the miRNeasy micro kit (Qiagen) according to the manufacturer’s instructions. RNAs, 500ng, were reversed-transcribed using the High Capacity RNA-to-cDNA Kit (Applied Biosystems). Real-time qPCR was performed in a Step-One Plus system (Applied Biosystems) using All-in-One qPCR Mix (GeneCopoeia). Quantitative PCR consisted of 40 cycles, 15 s at 95°C and 30 s at 60°C each, followed by dissociation curve analysis. The ΔCt was calculated by subtracting the Ct for the endogenous control gene GAPDH from the Ct of the gene of interest. Mouse primer sequences are listed in Table 3. Relative quantification was performed using the ΔΔCt method^4^ and expressed as a -fold change relative to control by calculating 2^-ΔΔCt^.

### Human induced pluripotent stem cell-derived NSC culture

Human induced pluripotent stem cells (iPSCs) were differentiated into prepatterned Activin A-treated neural stem cells (NSCs) using the following protocol. Briefly, iPSC colonies were detached using 1 mg/ml collagenase (Type IV, Thermo Fisher Scientific, 17104019) in Gibco KnockOut DMEM/F-12 medium (Thermo Fisher Scientific, 12660012), and the resulting cell clumps were transferred to a 0.1% agarose (Sigma-Aldrich, A9414) coated low-attachment petri dish in embryonic stem (ES) culture medium [Gibco KnockOut DMEM/F12 supplemented with 20% Gibco KnockOut Serum Replacement (Thermo Fisher Scientific, 10828028), 2.5 mM L-glutamine (Thermo Fisher Scientific, 25030081), 1 X Non-Essential Amino Acids (NEAAs) (Thermo Fisher Scientific, 11140050), 15 mM HEPES (Thermo Fisher Scientific, 15630106), 0.1 mM β-mercaptoethanol (Thermo Fisher Scientific, 31350010), 100 U/ml penicillin-Streptomycin (Thermo Fisher Scientific, 15140122). Every 2 d, 25% of ES medium was replaced by embryoid body (EB) differentiation medium [DMEM (Corning, 10-013-CV) supplemented with 20% FBS (Thermo Fisher Scientific, 16000036), 1 X NEAA, 2 mM L-glutamine, 100 U/ml penicillin-Streptomycin]. At day 8, 100% of the culture medium was EB medium. At day 10, the embryoid bodies were attached to dishes coated with poly-L-ornithine (1:1000 in PBS; Sigma-Aldrich, P3655,) and laminin (1:100 in KnockOut DMEM/F-12; Sigma-Aldrich, L2020,), and cultured in neural induction medium [DMEM/F12 supplemented with 1 X N2 (Thermo Fisher Scientific, 17502001), 100 U/ml penicillin-Streptomycin] and 25 ng/ml βFGF (Peprotech, 100-18B) and 25 ng/ml Activin A (Peprotech, 120-14P). Media change was performed every 2 d. Rosettes were harvested after 7–10 d, plated on poly-L-ornithine- and laminin-coated dishes, and cultured in Neural Proliferation Medium [NPM; Neurobasal medium (Thermo Fisher Scientific, 21103049), B27-supplement 1 X (Thermo Fisher Scientific, 17504001), GlutaMAX 1 X (Thermo Fisher Scientific, 35050061), 10 ng/ml leukemia inhibitory factor (Peprotech, 300-05), 100 U/ml penicillin-Streptomycin] supplemented with 25 ng/ml β-FGF and 25 ng/ml Activin A. The resulting NSCs were passaged and maintained in this same medium. These prepatterned Activin A-treated NSCs were validated by immunofluorescence analysis and labeled positively for putative NSC markers, namely Nestin, SOX1, SOX2, and PAX6.

### Adenovirus transduction of human NSCs

For the ADV transduction experiments human iPSC derived NSCs were plated at 700,000 cells per well of a 6-well plate in 2 ml NPM supplemented with 25 ng/ml βFGF (Peprotech, 100-18B) and 25 ng/ml Activin A (Peprotech, 120-14P). Two day after plating, cells were transduced at MOI of 10 with ADV-CMV-*FOXF2*-mCherry (SignaGen Laboratories, SL100701), ADV-CMV-OLIG2-mCherry (SignaGen Laboratories, SL100756) and ADV-CMV-STAT1-6xHN (SignaGen Laboratories, SL110858) and MOI of 0.75 with ADV-CMV-EGFP (SignaGen Laboratories, SL100708) in 1 ml NPM without penicillin/streptomycin antibiotic. Then, the cells were placed on an orbital shaker in 37°C for 1h. A complete media change was performed 24 h post-transduction. Non-transduced NSCs and NSCs transduced with ADV-EGFP were used as controls. Cells were harvested 4 days after transduction for gene expression and immunolabeling assays. ADV transduction in a human iPSC-derived NSC culture was performed twice, with three replicates each.

### Cell immunofluorescence of human NSCs

Cells were fixed using 4% paraformaldehyde in 0.1 M phosphate-Buffer Saline (PBS), pH 7.4 (Corning, 21-040-CV) for 30 min. Following three washes in phosphate-buffered saline, cells were permeabilized and blocked for 1 h at RT using 0.1% Triton X-100 (Thermo Fisher Scientific, 28313) and 4% donkey serum in PBS. Primary antibodies were added in the presence of blocking buffer overnight at 4°C. Secondary antibodies (1:500) were added following three PBS washes in blocking buffer at RT for 1 h. The following primary antibodies were used for the immunofluorescence studies: rabbit anti-DARPP-32 (Santa Cruz, sc-11365, 1 :100) and rabbit anti-Opioid Receptor-Mu (Millipore, AB5511, 1:500). The secondary antibodies were donkey anti-rabbit IgG conjugated with Alexa-488 (Invitrogen, A12379) or Alexa-647 (Invitrogen, A22287). Images were acquired using a Biotek Cytation 5 microscope and were prepared using Fiji software (ImageJ)

### Quantitative real-time PCR of human NSCs

For qRT-PCR analysis of prepatterned Activin A-treated human NSCs, total RNA was isolated using the ISOLATE II RNA Mini Kit (Bioline, BIO-52072). cDNA was prepared from 300 ng of RNA in a total reaction volume of 20 µl using the Sensi-FAST cDNA synthesis kit (Bioline, BIO-65053). RT-PCR reactions were set up in a 384-well format using 2X SensiFAST Probe No-ROX Kit (Bioline, BIO-86005) and 1 µl of cDNA per reaction in a total volume of 10 µl. RT-PCR was performed on the Roche LightCycler 480 instrument. Quantitative PCR consisted of 40 cycles, 15 s at 95°C and 30 s at 60°C each, followed by dissociation curve analysis. The ΔCt was calculated by subtracting the Ct for the endogenous control gene β-actin from the Ct of the gene of interest. Human primer sequences are listed in **Table 4**. Relative quantification was performed using the ΔCt method and is expressed as a fold change relative to control by calculating 2^-ΔCt^.

**Table 4.**
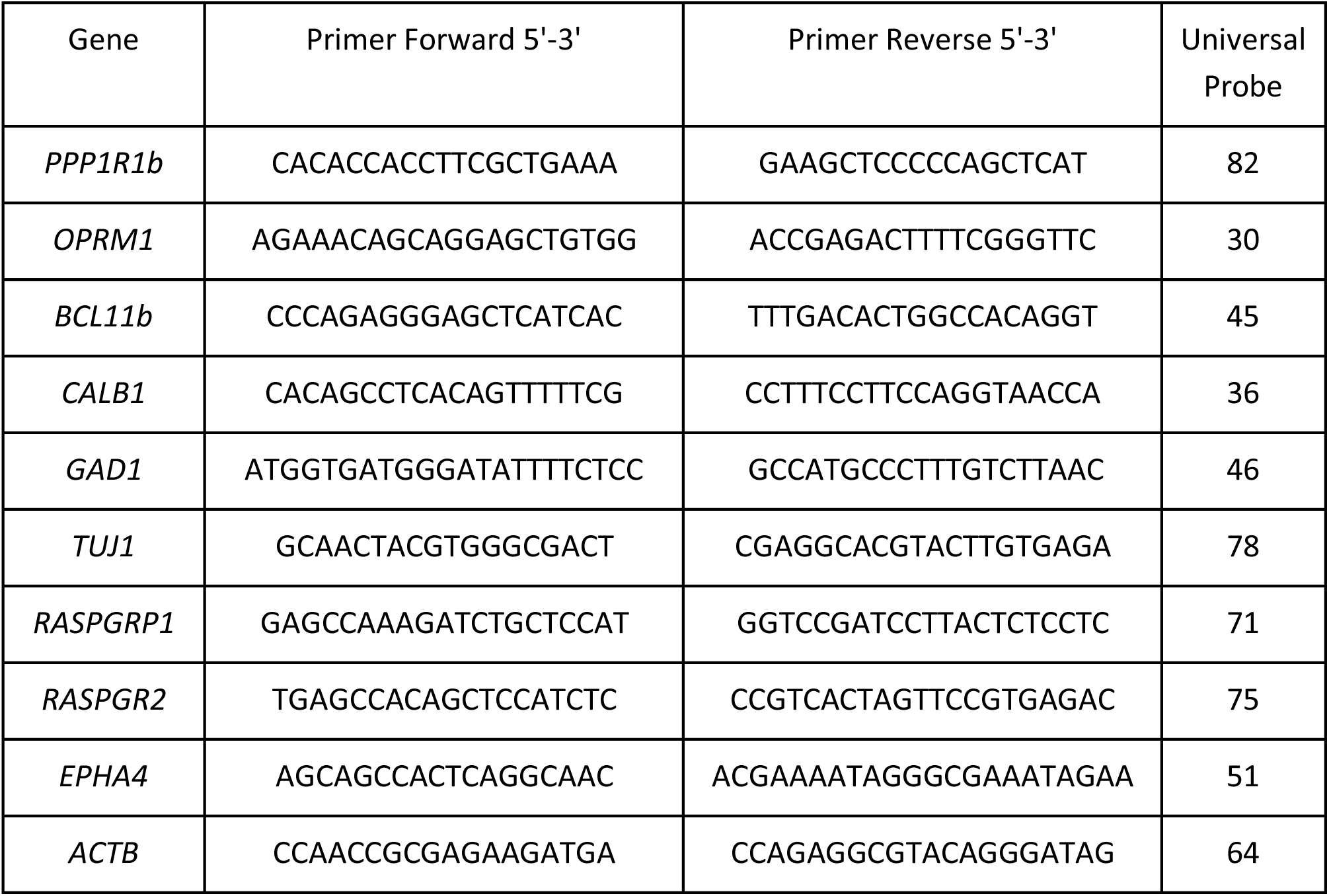
qRT-PCR human primer sequences.

### Generation of RNA-seq libraries

For RNA-seq, EGFP positive and negative cells were sorted into low-binding tubes containing Arcturus PicoPure Extraction buffer. RNA was isolated in accordance with the PicoPure RNA Isolation kit manufacturer’s instructions, which included a DNase treatment step. Samples were eluted in RNase-free water and stored at -80oC until preparation of RNA-Sequencing libraries using the Takara Clontech Laboratories SMARTer Stranded Total RNA-Seq Pico Kit, according to the manufacturer’s instructions. Following construction of the RNA-seq libraries, libraries were analyzed on an Agilent High Sensitivity D1000 TapeStation, and quantification of the libraries was performed using the KAPA Library Quantification Kit.

### RNA-seq

RNA-seq was carried out at New York University using Illumina HiSeq 4000 Paired-End 150 Cycle Lane from purified RNA (PicoPurePicoPure® RNA Isolation Kit KIT0204 Arcturus (ThermoScientific) and SMARTer Stranded Total RNA-Seq Kit - Pico Input Mammalian (250 pg–10 ng RNA) (635005, Clontech). The low-quality base (quality score lower than 20), as well as the adapters of the raw reads from the sequencing experiments, were removed using Trim Galore! (https://www.bioinformatics.babraham.ac.uk/projects/trim_galore/). The external and internal rRNA contamination were filtered through SortMeRNA 2.1b^116^. Then the filtered raw reads were then mapped to the Genome Reference Consortium Mouse Build 38 striosome release 6(GRCm38.p6) assembly by GENCODE using STAR 2.7.2b^117^. The counts of reads mapped to known genes were summarized by featureCounts, using GENECODE release M22 annotation (GSE143276). The data discussed in this publication have been deposited in NCBI’s Gene Expression Omnibus GEO Series accession number, GSE143276.

Next, R Bioconductor package DESeq2^118^ was used to normalize raw read counts logarithmically and perform differential expression analysis. Differentially expressed genes were based on an arbitrary cutoff of adjusted p-value less than 0.01 (Table 1). We found the regulation pattern of a gene with the same EGFP status is more likely to be the same across replicates, whereas it is more likely to be different when the EGFP status is different between replicates with the exception of *Xist. Xist* is a gene highly expressed in females only. The regulation pattern of *Xist* is consistent with gender differences of the samples confirming the identity of each sample (Fig. 1e).

### Terminology enrichment analysis and pathway enrichment analysis

Enrichment analysis was performed on gene clusters in specific databases to determine whether a specific biological annotation could be considered as significantly represented under the experiment result. Both terminology enrichment analysis and pathway enrichment analysis were conducted by clusterProfiler^119^, a Bioconductor package. In our analysis, biological process (BP), molecular function (MF), and cellular component (CC) terms in gene ontology (GO)^57^ as well as pathway annotations derived from Kyoto Encyclopedia of Genes and Genomes (KEGG) were chosen to identify predominant biological processes of the differentially expressed gene clusters and differentially expressed transcription factor clusters involved in the development of the striosome neurons. We conducted both analyses on the differentially expressed gene clusters with the arbitrary cutoff of adjusted p-value less than 0.01 and the absolute Log2 fold change greater than 0, 1, and 2 respectively, and we conducted both analyses on the differentially expressed transcription factor clusters with the arbitrary cutoff of adjusted p-value less than 0.01 and the absolute Log2 fold change greater than 0 and 1 respectively (Table 3).

### Transcription factor enrichment analysis and co-expressor enrichment analysis

Both transcription factor enrichment analysis and co-expressor enrichment analysis were conducted by Enrichr^54^, a comprehensive online tool for doing enrichment analysis with a variety of biologically meaningful gene set libraries (Table 3). In our analysis, ChEA^120^ and ENCODE^121^ databases were chosen to identify the significant upstream transcription factors regulating genes and other TFs differentially expressed in striosome cells and matrix cells respectively and ARCHS4 database was chosen to identify the significant co-expressors of those differentially expressed genes and transcription factors. An arbitrary cutoff of adjusted p-value less than 0.01 and the absolute log2 fold change greater than 0 and 1 were chosen (Table 1).

### GeneMANIA gene regulatory network analysis

GeneMANIA^55^ is an online tool using published and computational predicted functional interaction data among proteins and genes to extend and annotate the submitted gene list by their interactive biological pathways and visualize its inferred interaction network accordingly. We used GeneMANIA to conduct the interaction network inference analysis to transcription factors enriched in either the striosome or matrix compartments with the arbitrary cutoff of adjusted p-value less than 0.01 and the absolute log2 fold change greater than 1.

### Gene regulatory network inference through data curation

A gene regulatory network links transcription factors to their target genes and represents a map of transcriptional regulation. We used all the transcription factors and their target gene data curated by ORegAnno^68^ to build the network. In order to simplify the network, we only chose the compartmental differentially expressed transcription factors that are high on the hierarchy. In other words, only the differentially expressed transcription factors that served as a regulator of other differentially expressed transcription factors were chosen as the candidates of our gene regulatory network.

### Gene Set Enrichment Analysis

GO^122^ was performed using ranked list of differential gene expression with parameters set to 2000 gene-set permutations and gene-set size between 15 and 200. The gene-sets included for the GSEA analyses were obtained from Gene Ontology (GO) database (GOBP_AllPathways), updated September 01, 2019 (http://download.baderlab.org/EM_Genesets/). An enrichment map (version 3.2.1 of Enrichment Map software^123^) was generated using Cytoscape 3.7.2 using significantly enriched gene-sets with an FDR <0.05. Similarity between gene-sets was filtered by Jaccard plus overlap combined coefficient (0.375). The resulting enrichment map was further annotated using the AutoAnnotate Cytoscape App.

### Statistical comparison with HD mouse transcriptomes

Fisher’s exact tests^124^ were conducted to identify overlap between differentially expressed gene sets from similar experiments. We performed this by filtering all datasets on GEO database based on cell type, the organism, and the type of sequencing methods. The query was limited to all datasets relevant to HD research in mouse and human and high throughput genomics experiments which would include up to 84 GEO entries. Among these 84 entries, thirteen of them provided differentially expressed gene lists for further meta-analysis. Two of the studies had derived two differentially expressed gene lists from two independent experiments. Fourteen publicly accessible neurodegenerative disease datasets (GSE9038^125^, GSE19780^126^, GSE65774^72^, GSE78274^72^, GSE89505^127^, GSE59571^128^, GSE139847^129^, GSE48962^130^, GSE129473^131^, GSE104091^132^, GSE74201^71^, GSE79666^133^, GSE95344^69^, GSE49036^134^) were used to do the comparative analysis. We used a background gene dataset 55376 for mouse data, in the GENECODE release M22 genome annotation data^121^, and a background gene dataset 58219 for human data, in the GENECODE release 25 genome annotation data. We considered the expression pattern of the gene list in the test. We conducted two experiments using the Fishers’ exact test. For the first test, only the genes that share the same expression pattern with patch cell are counted as overlapping genes. In this test, since we treated the matrix cells as a control, only the differentially expressed genes that share the same pattern of expression (enriched in striosome or matrix) and are in direct correlation in both datasets were counted as overlapping genes. For the second test, we compare only the genes that share the same expression pattern with matrix cells. In other words, only the genes that are differentially expressed in both datasets and their pattern of expression are in inverse correlation in both datasets were counted as overlapping genes in this test.

### Data processing

The preprocessing of ATAC-seq data involved the following steps:

- <u>Alignment.</u> Sequencing reads were provided by the sequencing center demuxed and with adaptors trimmed. Reads from each sample were aligned on GRCh38-mm10 reference genome using the STAR aligner^117^ (v2.5.0) with the following parameters:

*--alignIntronMax 1*
*--outFilterMismatchNmax 100*
*--alignEndsType EndToEnd*
*--outFilterScoreMinOverLread 0.3*
*--outFilterMatchNminOverLread 0.3* This produced a coordinate-sorted BAM file of mapped paired-end reads for each sample. We excluded reads that: [1] mapped to more than one locus using SAMtools^135^; [2] were duplicated using PICARD (v2.2.4; http://broadinstitute.github.io/picard); and [3] mapped to the mitochondrial genome.
- <u>Quality control metrics.</u> The following quality control metrics were calculated for each sample: [1] total number of initial reads; [2] number of uniquely mapped reads; [3] fraction of reads that were uniquely mapped and additional metrics from the STAR aligner; [4] Picard duplication and insert metrics; [5] normalized strand cross-correlation coefficient (NSC) and relative strand cross-correlation coefficient (RSC), which are metrics that use cross-correlation of stranded read density profiles to measure enrichment independently of peak calling. Supplementary Table 1 describes the main QC metrics. The bigWig tracks for each sample were manually inspected. None of the libraries failed QC and visual inspection and 8 libraries were subjected to further analysis.
- <u>Selection and further processing of samples meeting quality control.</u> We subsequently subsampled samples to a uniform depth of 10 million paired-end reads and merged the BAM-files of samples from the same cell type. We called peaks using the Model-based Analysis of ChIP-Seq (MACS)^**136**^ v2.1 (https://github.com/taoliu/MACS/). It models the shift size of tags and models local biases in sequencability and mapability through a dynamic Poisson background model. We used the following parameters:

*--keep-dup all*
*--shift -100*
*--extsize 200*
*--nomodel* We created a joint set of peaks requiring each peak to be called in at least one of the merged BAM-files. That is, if a peak was identified in just one or more samples it was included in the consensus set of peaks. If two or more peaks partially overlapped, the consensus peak was the union of bases covered by the partially overlapping peaks. After removing peaks overlapping the blacklisted genomic regions, 69,229 peaks remained. We subsequently quantified read counts of all the individual non-merged samples within these peaks, again, using the feature counts function in RSubread^137^ (v.1.15.0). We counted fragments (defined from paired-end reads), instead of individual reads, that overlapped with the final consensus set of peaks. This resulted in a sample by peak matrix of read counts, obtained using the following parameters:

> *allowMultiOverlap = F,*
>
> *isPairedEnd = T,*
>
> *strandSpecific = 0,*
>
> *requireBothEndsMapped = F,*
>
> *minFragLength = 0,*
>
> *maxFragLength = 2000,*
>
> *checkFragLength = T,*
>
> *countMultiMappingReads = F,*
>
> *countChimericFragments = F*

### Differential analysis of chromatin accessibility

To identify genomic regions with significant regional differences in chromatin structure among the two cell types, we performed a statistical analysis of chromatin accessibility. Here, chromatin accessibility is assessed by how many ATAC-seq reads overlap a given OCR: the higher the read count, the more open the chromatin is at a given OCR. For this, we performed the following steps:

- <u>Read counts:</u> As a starting point, we used the sample-by-OCR read count matrix described in the previous section (8 samples by 69,229 OCRs). From here, we subsequently removed 9 OCRs using a filtering of 0.5 CPM in at least 50% of the samples, resulting in our final sample-by-OCR read count matrix (8 samples by 69,220 OCRs). Next, we normalized the read counts using the trimmed mean of M-values (TMM) method^138^.
- <u>Covariate exploration:</u> To explore factors affecting the observed read counts we examined several biological and technical sample-level variables. For these covariates (e.g. number of peaks called in the sample, chrM metrics, RSC and NSC, and Picard insert metrics) we normalized to the median of the cell. We next assessed the correlation of all the covariates with the chromatin accessibility values in the normalized read count matrix to determine which of these variables should be candidates for inclusion as covariates in the differential analysis. We did this using a principal component analysis of the normalized read count matrix and by examining which variables were significantly correlated with the high-variance components (explaining > 1% of the variance) of the data. We did not identify any significant association even when we used a lenient false discovery rate threshold of 0.2.
- <u>Differential analysis:</u> We used the edgeR package138. to model the normalized read counts using negative binomial (NB) distributions. The estimateDisp function was used to estimate an abundance-dependent trend for the NB dispersions^139^. To normalize for compositional biases, the effective library size for each sample was estimated using the TMM approach as described above. For each open chromatin region, we applied the following model for the effect on chromatin accessibility of each variable on the right-hand side:

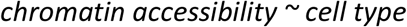 Then, for each OCR, the cell type coefficient was statistically tested for being non-vanishing. A quasi-likelihood (QL) F-test was conducted for each OCR using the glmQLFTest function^140^ from the edgeR package, with robust estimation of the prior degrees of freedom. p-values were then adjusted for multiple hypothesis testing using false discovery rate (FDR) estimation, and the differentially accessible regions of chromatin were determined as those with an estimated FDR below, or at, 5%.

### Annotation of OCRs

We used the gene annotations form the org.Mm.eg.db (version 3.8.2) package for all analyses in this paper. We assigned the closest gene and the genomic context of an ATAC-seq OCR using ChIPSeeker^141^. The genomic context was defined as promoter (+/-3Kb of any TSS), 5’-UTR, 3’-UTR, exon, intron, distal intergenic and downstream.

Transcription factor binding motif analysis of ATAC-seq data was performed using HOMER suit function findMotifsGenome.pl tool. Only known motifs from HOMER’s motif database were considered.

## Figure Legends

**Table 1**. Differential expressed genes and bioinformatics analysis of transcriptions of FACs EGFP^+^ and EGFP^−^ cells populations from the striata of PND3 *Nr4a1*-GFP mice.

**Table 2**. Transcriptional regulation of striosome and matrix specific transcription networks overlaps with dysregulated transcriptomics in neurological diseases.

## Supplementary Figure Legends

**Figure S1.**
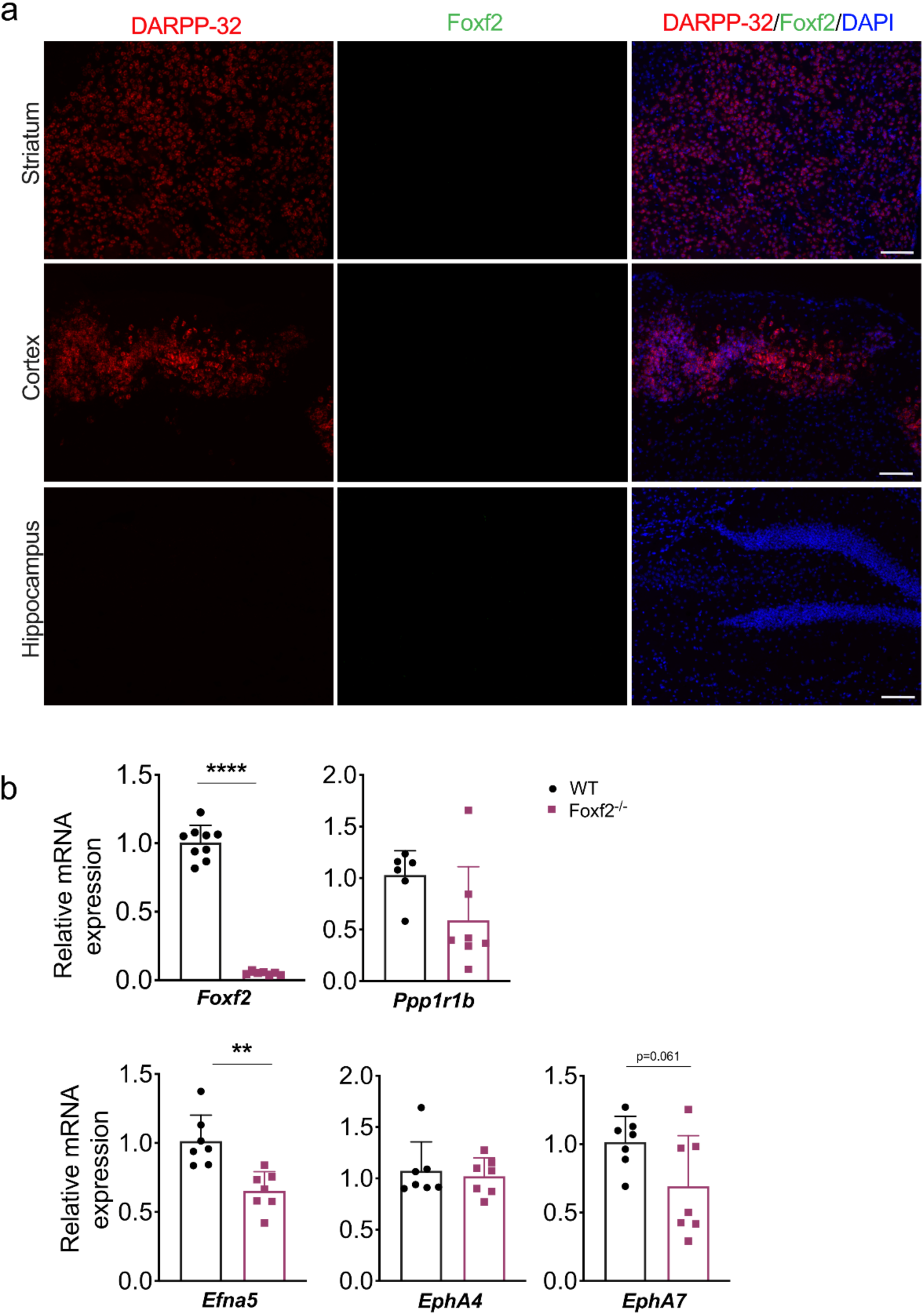
*Foxf2* mRNA is undetectable in adult striatum and Ppp1r1b mRNA levels are almost normal in E18.5 *Foxf2*-null striatum. **a)** Low power view of RNAscope assay visualization of *Foxf2* and *Ppp1r1b*/DARPP-32 mRNA in adult mouse striatum, cortex and hippocampus indicating that *Foxf2* is not expressed in neurons in adult brain. **b)** RT-qPCR from striatal RNA from E18.5 *Foxf2* WT littermates indicating that *Foxf2* expression is abolished in the null mouse and that the level of *Ppp1r1b or Epha4* mRNA is not significantly altered by *Foxf2* deletion unlike *Efna5* mRNA. n=7 for *Foxf2*^-/-^ and n=9 for WT. Unpaired t-test **p<0.001, ****p<0.0001. Error bars represent standard deviation.

**Figure S2.**
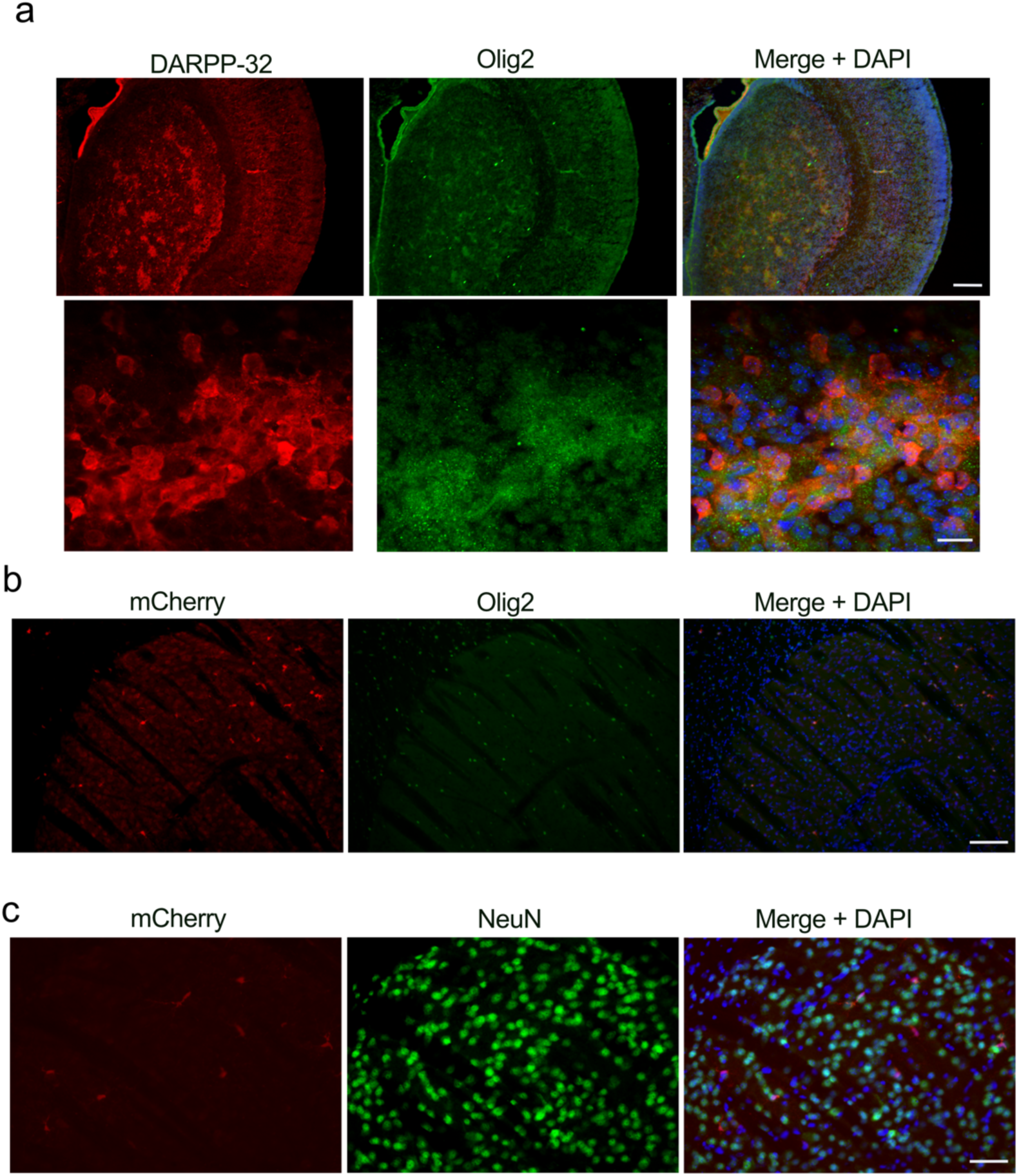
Confirmation of striosome localization of Olig2 and analysis of Olig2 OCR-driven expression in adult founders. **a)** Low and high-power magnification of Olig2 (green), DARPP-32 (red) and DAPI immunolabelling on WT PND3 coronal sections (16 1m) using a second antibody against Olig2 (gift Dr. Wichterle)^115^ showing the localization of Olig2 within the DARPP-32 positive striosomes. Scale bars correspond to 200 µm and 20 µm respectively. **b)** Representative low power micrographs of mCherry (red), Olig2 (green) and DAPI immunolabelling on coronal sections (30 1m) from adult *Olig2* -OCR transgenics indicating co-localization with Olig2. Scale bar: 200 µm. **c)** Representative micrograph of mCherry (red), NeuN (green) and DAPI immunolabelling on coronal sections (30 1m) from adult *Olig2* -OCR transgenics indicating absence of striatal neuronal expression. Scale bar: 50 µm

**Fig. S3.**
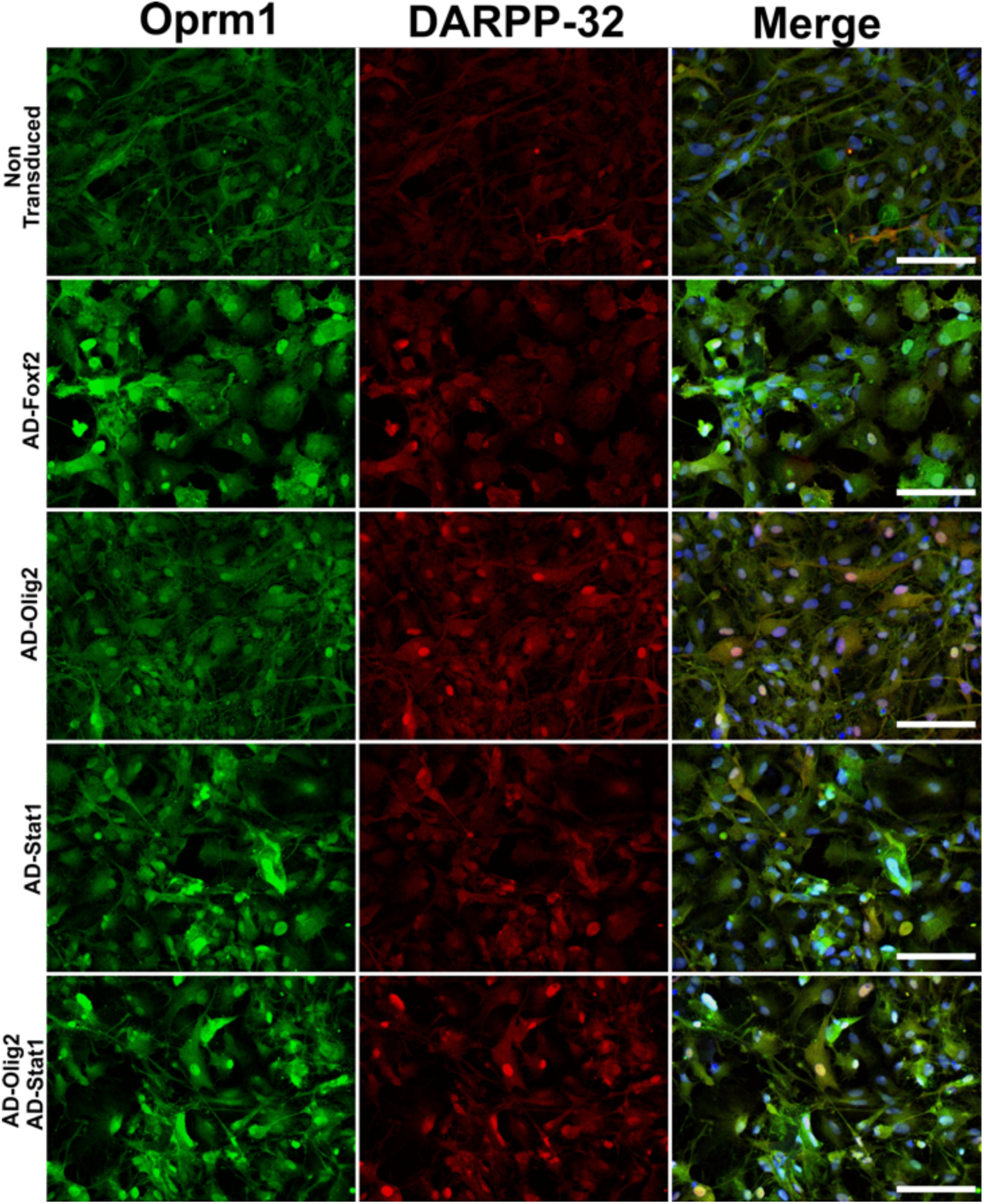
Foxf2, Olig2, and Stat1, alone and in combination, promote MSN differentiation in NSCs from human isogenic control C116 induced pluripotent stem cells. **a)** C116-NSCs transduced for 4 days with ADV-*FOXF2*, ADV-*OLIG2* and ADV-*STAT1* were immunostained with Oprm1 (green) and DARPP-32 (red). Non-transduced cells and AD-GFP were used as control. Scale bars: 100 µm.

**Supplementary Table 1**. Uniquely mapped reads after removing duplicate reads 322 million (average of 40.3 million) and those aligning to the mitochondrial genome.

**Supplementary Table 2**. Comparison of striosome vs. matrix peaks identified ∼44% of OCRs that were significant after multiple testing corrections.

## Acknowdedgements

This research was supported by the Department of Health and Human Services|National Institutes of Health|National Institute of Neurological Disorders and Stroke Grant R01-NS-100529 and the Collaborative Center for X-linked Dystonia Parkinsonism (to L.M.E. and M.E.E.). The Taube HD Stem Cell Consortium also provided support (L.M.E.) as well as NLM fellowship grant T15LM007442 (H.B). A postdoctoral fellowship was provided to K.T.T. from the Collaborative Center for X-linked Dystonia Parkinsonism.

## Author contributions

Conceptualization: L.M.E. and M.E.E. Methodology: M.D.C., S.S., K.T.T., C.C., J.M.,C.G.A.,H.B., J.B., P.A., J.F.F., P.C., P.R., S.D.M., L.M.E. and M.E.E. Formal analysis: M.D.C., S.S., K.T.T., C.G.A., H.B., J.F.F. P.R., S.D.M. Investigation: M.D.C., S.S., K.T.T., C.C., J.M.,C.G.A.,H.B., J.B., P.A., J.F.F., P.C., P.R., S.D.M., L.M.E. and M.E.E. Resources P.C., P.C. L.M.E. M.E.E. Writing (original draft) M.D.C., K.T.T. S.S., P.R.,L.M.E., M.E.E. Writing (reviewing and editing) M.D.C., S.D.M., S.S., P.R.,L.M.E., M.E.E. Visualization M.D.C., S.S., K.T.T., C.G.A., L.M.E., M.E.E. Supervision: L.M.E. and M.E.E. Project adminstration and research funding: L.M.E. and M.E.E.

## Additional information

Data availability statement: all the raw and normalized count are in the GEO data depository (accession codes: GEO is GSE143727 for ATAC-seq and GSE143276 for the RNAseq). Figure 1, 4-8 have associated raw data. All the data will be available a year from the date of the manuscript publication. Code availability statement: The codes used during the generation of the data is available in public depository Github that will be available after the publication of the manuscript.

